# A paralog-specific role of the COPI pathway in the neuronal differentiation of murine pluripotent cells

**DOI:** 10.1101/2020.01.27.921924

**Authors:** Manu Goyal, Xiyan Zhao, Mariya Bozhinova, Karla Lisette Andrade López, Cecilia de Heus, Sandra Schulze-Dramac, Michaela Müller-McNicoll, Judith Klumperman, Julien Béthune

**Affiliations:** Junior Research Group, Cluster of Excellence CellNetworks, D-69120 Heidelberg, Germany; Heidelberg University Biochemistry Center, D-69120 Heidelberg, Germany; Section Cell Biology, Center for Molecular Medicine, University Medical Center Utrecht, Utrecht University, Utrecht 3584 CX, The Netherlands; RNA Regulation Group, Cluster of Excellence ‘Macromolecular Complexes’, Institute of Cell Biology and Neuroscience, Goethe University Frankfurt, Frankfurt/Main, Germany

## Abstract

Coat protein complex I (COPI)-coated vesicles mediate membrane trafficking between Golgi cisternae as well as retrieval of proteins from the Golgi to the endoplasmic reticulum. There are several flavors of the COPI coat defined by paralogous subunits of the protein complex coatomer. However, whether paralogous COPI proteins have specific functions is currently unknown. Here we show that the paralogous coatomer subunits γ1-COP and γ2-COP are differentially expressed during the neuronal differentiation of mouse pluripotent cells. Moreover, through a combination of genome editing experiments, we demonstrate that whereas γ-COP paralogs are largely functionally redundant, γ1-COP specifically promotes neurite outgrowth. Our work stresses a role of the COPI pathway in neuronal polarization and provides evidence for distinct functions for coatomer paralogous subunits in this process.

## INTRODUCTION

In eukaryotes, membrane trafficking of cargo proteins and lipids is crucial to maintain cellular homeostasis and intracellular organelle identity. In the early secretory pathway in mammals, coat protein complex II (COPII) vesicles mediate the export of cargo from the endoplasmic reticulum (ER) to the Golgi apparatus whereas COPI vesicles promote the retrieval of proteins from the Golgi to the ER, and intra-Golgi transport. COPI vesicles are formed at the Golgi through the GTP-dependent recruitment of a coat protein complex, termed coatomer, by the small GTPase ARF1. Once recruited to membranes, coatomer polymerizes to form a lattice that shapes a nascent bud that eventually pinches off as a small-coated vesicle^1^. Through sorting signals exposed on their cytoplasmic domains, transmembrane cargo proteins interact with coatomer and are taken up in COPI vesicles^2^. Coatomer is made of seven equimolar COP subunits (α-,β-,β′-,γ-,δ-,ε- and ζ-COP) that are highly conserved from yeast to human. In mammals, two coatomer subunits come as two paralogs: γ1-COP and γ2-COP that share 80% protein sequence identity and that are encoded by the *Copg1* and *Copg2* genes^3^, and ζ1-COP and ζ2-COP encoded by the *Copz1* and *Copz2* genes^4^. In the COPII system, paralogs of the SEC24 subunit expand the cargo repertoire of COPII vesicles by providing distinct binding sites for specific sorting motifs^5–9^. By contrast, proteomics profiling of paralog-specific COPI vesicles generated *in vitro* from HeLa cells revealed no major difference in their cargo content^5^ and to date no specialized function has been ascribed to the paralogous COP subunits, which until now have thus been considered as functionally redundant.

Whereas the general mechanisms of cargo sorting and formation of COPI vesicles are well described, cell type-specific functions are much less well studied. Several lines of evidence, however, suggest tissue-specific functions of the otherwise essential COPI pathway. Indeed, mutations affecting COP subunits have been associated to diseases, notably neurodegenerative disorders^10–13^. Moreover, binding of α-COP to the survival motor neuron (SMN) protein seems to promote neurite outgrowth in motor neurons^14^. As defects in the COPI pathway lead to specific effects in the nervous system, we decided to analyze the expression profile and function of the γ-COP paralogs during neurogenesis. By examining publicly available mRNA expression profiling data we found that *Copg1*, but not *Copg2*, is upregulated as mouse embryonic stem cells (mES) differentiate into neurons. We observed the same in murine pluripotent P19 cells, another model cell line for neuronal differentiation. Specific knockout (KO) of *Copg1* or *Copg2* in P19 cells revealed that whereas both gene disruptions led to slower cell growth, neither of the two paralogs alone is essential for cell viability. Remarkably, whereas KO of *Copg2* did not affect retinoic acid-mediated P19 cell neuronal differentiation, disruption of *Copg1* led to the formation of loose embryoid bodies (EBs) and to reduced neurite outgrowth. Overexpression of γ2-COP or knock-in of *Copg2* in the *Copg1* locus revealed that whereas higher expression of γ2-COP can compensate for the loss of γ1-COP for the formation of EBs, γ1-COP is required to promote neurite outgrowth. Altogether, our findings support an essential role of COPI vesicles during neurogenesis and reveal for the first time a paralog-specific function for a COPI protein subunit.

## RESULTS

### Differential expression of *Copg1* and *Copg2* during neuronal differentiation

To investigate a potential paralog-specific function of COP subunits during neurogenesis we first examined publicly available RNAseq expression profiling data performed on mES, their derived neuronal progenitors (NP) and terminal neurons (TN)^15^. In the three differentiation stages *Copz2* was marginally expressed compared to the other COP subunits with expression levels 10 to 40 time lower than those of *Copz1* (Supplementary Fig. 1). Interestingly, whereas all other COP subunit-coding genes were expressed at similar levels at all three differentiation stages, *Copg1* was strongly upregulated in TN (Supplementary Fig. 1) suggesting γ1-COP might exert a unique function during the biogenesis of neurons. To examine whether a differential expression of *Copg1* and *Copg2* is also observed in another neuronal differentiation system we then analyzed mRNA and protein levels of the two γ-COP paralogs in P19 embryonal carcinoma cells. Compared to mES, P19 cells are closer to epiblast-derived stem cells and are a well-established and robust model system for neuronal differentiation^16–18^. Similar to mES, P19 cells can be differentiated into neurons applying a two-step differentiation protocol (Fig. 1a). Cells are first incubated with retinoic acid (RA)-containing medium under non-adherent conditions to form EBs (day 2-4). EBs are then dissociated and the cells seeded under adherent conditions to form post-mitotic neurons (day 6-8). Following this procedure, cell lysates were collected at different stages of differentiation and analyzed by western blot. As expected, the pluripotency marker Oct-4 was only detectable at day 0 whereas the neuron-specific class III beta-tubulin (β III-tubulin or Tuj-1 antigen) is strongly upregulated as the cells progress to day 8 (Fig. 1b). The expression levels of different COP subunits were also assessed by western blot (Fig. 1b) using previously characterized specific antibodies against each of the γ-COP paralog^19^ and by RT-qPCR (Fig. 1c). To analyze the relative expression of the two γ-COP paralogs independent of antibody-dependent signal intensities, we considered the ratio of the γ1-COP over γ2-COP signals during differentiation. This ratio was similar in pluripotent cells and EBs, by contrast there was relatively more γ1-COP in neurons (Fig. 1d). This was corroborated at the mRNA levels as *Copg2* mRNA levels decreased at day 8 of differentiation compared to the transcripts of all the other COP subunits including *Copg1* (Fig. 1c, e). Altogether, the expression of γ1-COP is increased compared to γ2-COP during the late stage of neuronal differentiation of pluripotent cells.

**Figure 1:**
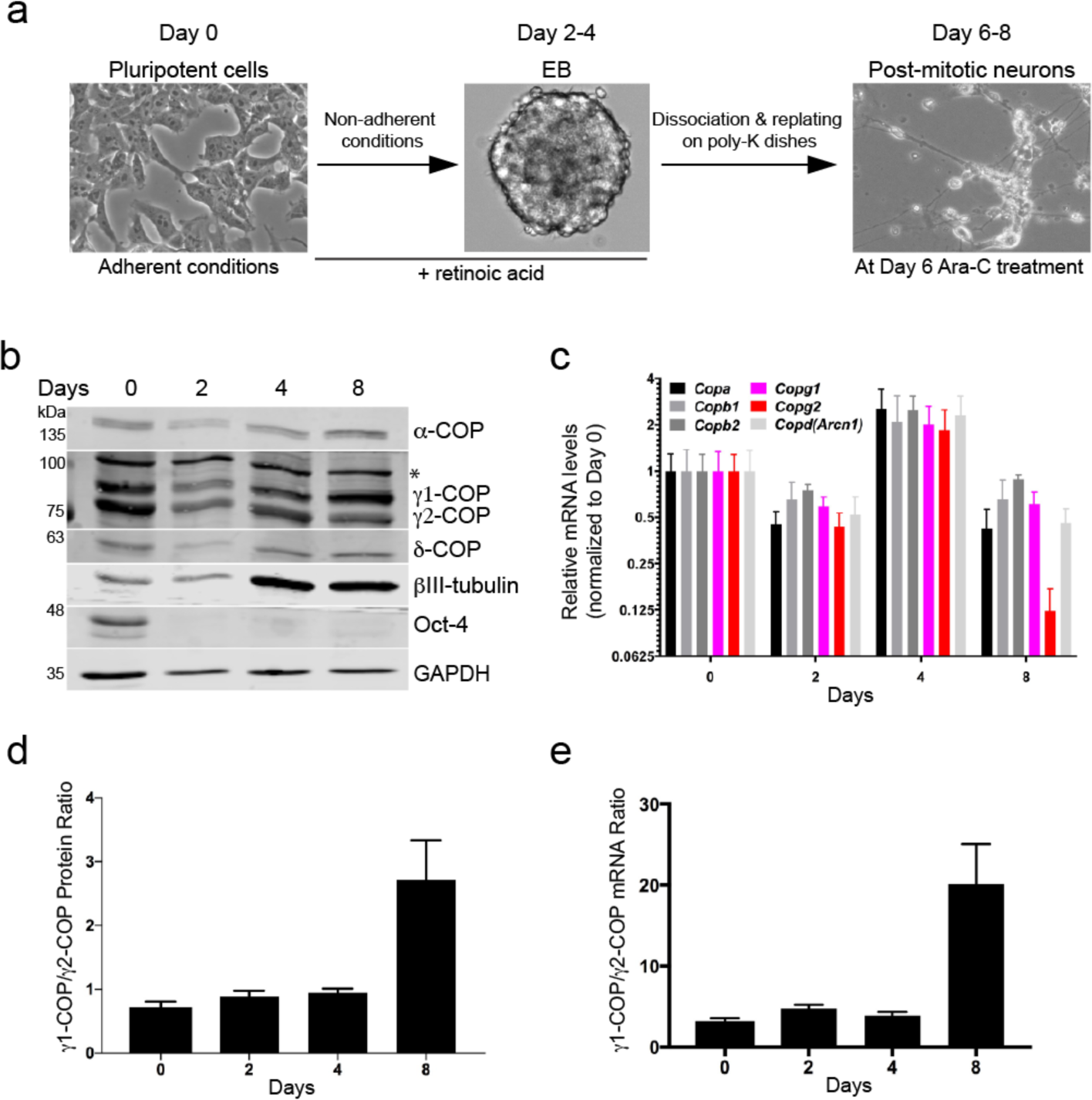
Differential expression of γ-COP paralogs during neuronal differentiation of P19 cells. (a) Schematic representation of the two-step differentiation protocol. (b) Western blot analysis of P19 cell lysates at different stages of differentiation. The asterisk (*) marks a non-specific signal. (c) Relative expression of mRNAs coding for the indicated COP subunits during neuronal differentiation by RT-qPCR. (d-e) Quantification (n = 3) of the γ1-COP/γ2-COP protein and mRNA ratios from (b) and (c) respectively.

### Disruption of *Copg1* or *Copg2* in P19 cells

To study potential paralog-specific functions of COPI vesicles we then generated *Copg1* and *Copg2* knockout (KO) P19 cells. To do so we followed a Cas9-mediated genome editing approach using individual specific sgRNAs directed against either *Copg1* or *Copg2* (see methods). After clonal selection, we obtained cell lines that showed absence of γ1-COP (*Copg1^-/-^* cells) or γ 2-COP (*Copg2^-/-^* cells) by western blot analysis (Fig. 2a). Moreover, bi-allelic disruption of *Copg1* or *Copg2* was confirmed by sequencing analysis of the respective genomic loci (Supplementary Fig. 2). Importantly, the knockout and clonal selection procedure did not lead to loss of expression of the pluripotency markers Oct-4 and NANOG (Fig. 2a), indicating that the KO cell lines are still undifferentiated cells.

**Figure 2:**
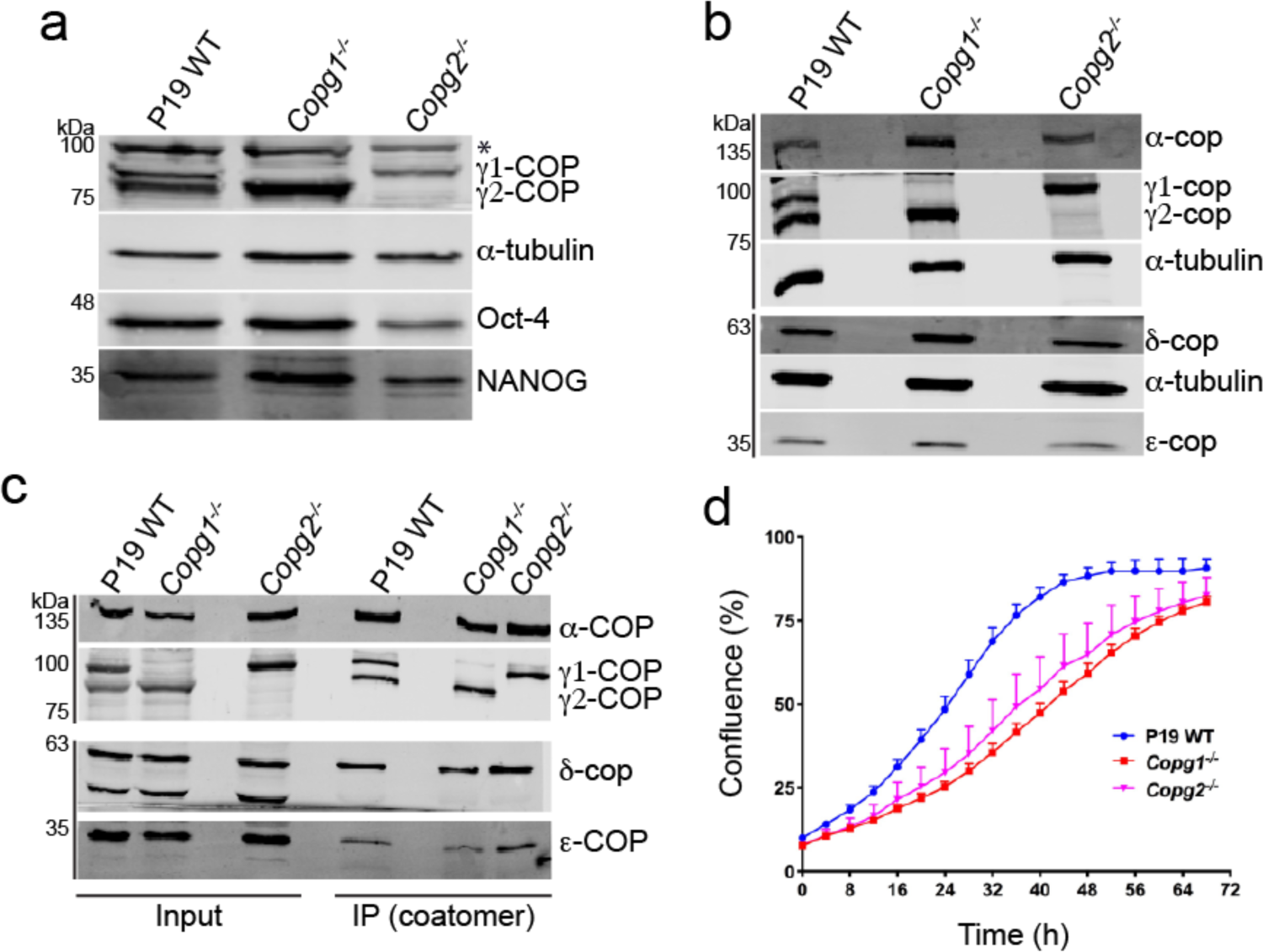
Characterization of P19 knockout cell lines. (a) Western blot analysis of the expression levels of γ1-COP and γ2-COP, the pluripotency markers Oct-4 and Nanog, and α-tubulin as a loading control in pluripotent P19 WT and KO cell lysates. The asterisk (*) marks a non-specific signal. (b) Western blot analysis of various COP subunits in pluripotent P19 WT and KO cell lysates (c) Immuno-precipitation of native coatomer from P19 WT or KO cell lysates using the CM1 monoclonal antibody. Various COP subunits were analyzed by western blotting as indicated. (d) Growth curves obtained from a real time proliferation assay in which the occupied area (% confluence) by P19 WT and KO cells was monitored over 72h. Curves were generated with the IncuCyte software (n=5, error bars are s.e.m.).

### Characterization of *Copg1* or *Copg2* KO in pluripotent P19 cells

As a functional COPI pathway is essential for viability, obtaining individual *Copg1* and *Copg2* knockout cells demonstrates that both γ-COP paralogs can fulfill essential functions of COPI vesicles and are thus at least partially redundant. It is hence expected that depletion of one or the other γ-COP paralog does not strongly affect the global expression levels of COP subunits and the assembly of the coatomer complex. To test these two predictions, we first analyzed the expression of several COP subunits in WT, and *Copg1* and *Copg2* KO cells. Western blot analysis revealed that expression of α-, δ- and ε-COP is similar in the three cell lines. As coatomer is an equimolar heptameric complex, the latter observation implies that in the KO cells the remaining γ-COP paralog is up-regulated. Accordingly, in western blot analyses, γ1-COP expression was reproducibly higher in *Copg2^-/-^* cells than in the WT situation, and conversely γ2-COP was up-regulated in *Copg1^-/-^* cells compared to WT (Fig. 2b, c). Hence, overexpression of one paralog compensates at least partially for the absence of the second one and thus ensures that global coatomer levels are relatively maintained. To verify whether coatomer subunits are correctly incorporated into the complex in the absence of one of the γ-COP paralogs, immunoprecipitation experiments from WT and KO cell lysates were performed using the CM1A10 (CM1) antibody, which specifically recognizes native assembled coatomer^20^ probably through its β′-COP subunit^21^. Comparable levels of α-, δ- and ε-COP were precipitated from all three cell lines indicating a correct assembly of coatomer in the absence of one specific γ-COP paralog (Fig. 2c). Finally, we performed a live cell proliferation assay by measuring the percentage surface occupancy of WT and KO cells in real time as they grow on a cell culture dish under standard conditions. In this assay, both *Copg1* and *Copg2* KO cells showed significantly slower proliferation rates than WT cells (Fig. 2d) suggesting that even though both γ-COP paralogs are individually sufficient to mediate essential COPI functions, paralog-specific non-essential functions exist.

### Aberrant Golgi morphology in *Copg1* and *Copg2* KO cells

As COPI vesicles are generated at the Golgi we then analyzed the morphology of that organelle by electron microscopy (EM) in WT, *Copg1^-/-^* and *Copg2^-/-^* cells (Fig. 3a). Quantitative assessment of 30 Golgi regions per cell line revealed an average higher number of Golgi stacks per cell in the absence of either γ1-COP or γ2-COP (2.7 Golgi/cell vs 1.8 in WT, Fig. 3b). In addition, the average area per Golgi stack was smaller in both KO cell lines (455-466 x10^3^ nm^2^ vs 730 x 10^3^ nm^2^ in WT, Fig. 3c). Specific to the depletion of γ1-COP was the observation of an increased number of Golgi peripheral vesicles (27.5% of observed Golgi for which vesicles accounted for more than 50% of the Golgi area vs. 13% and 12% WT and *Copg2^-/-^* cells respectively, Fig. 3d) indicative of increased Golgi fragmentation. Altogether, both γ1-COP and γ2-COP are necessary for correct assembly and/or maintenance of the Golgi structure.

**Figure 3:**
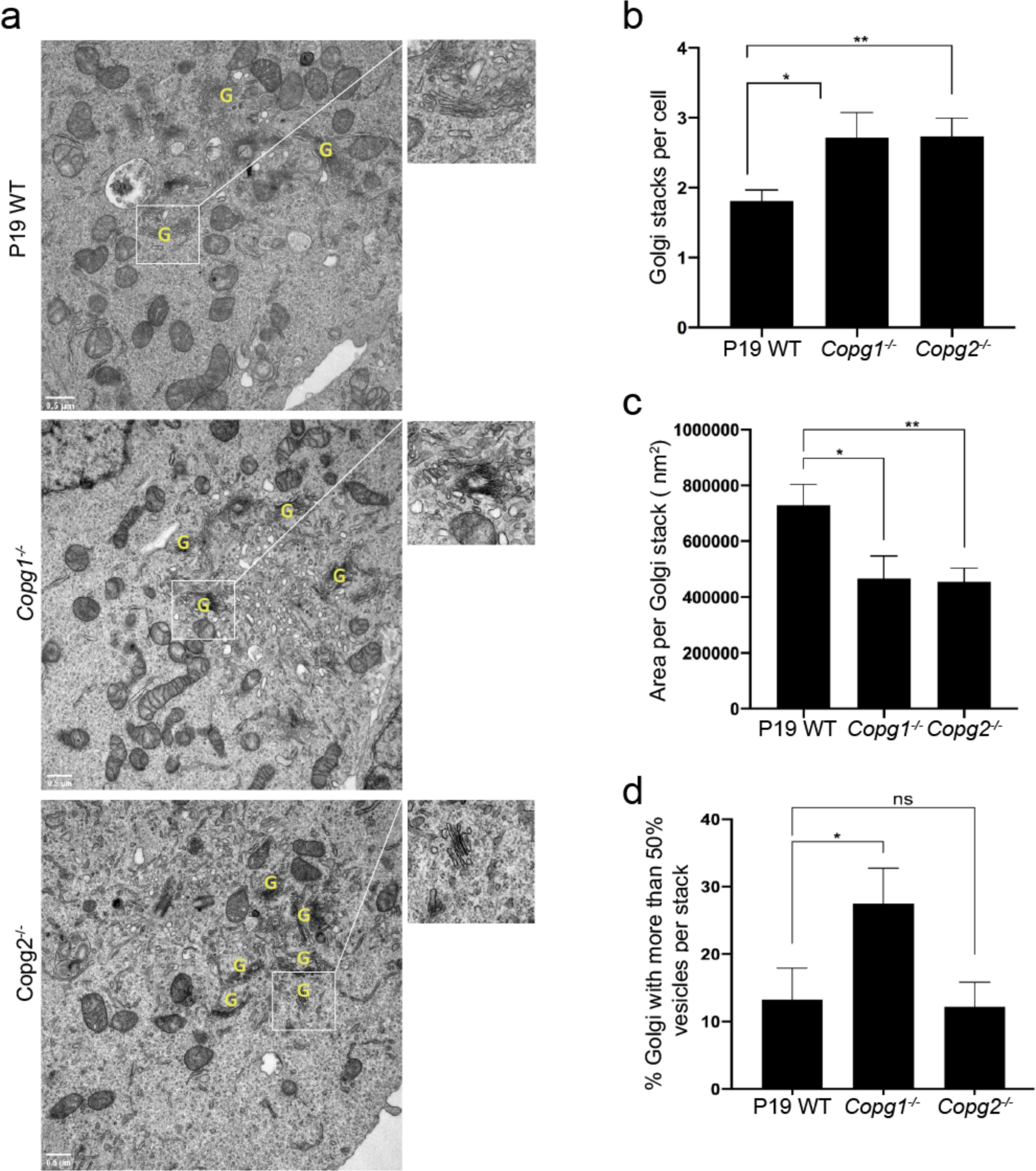
Depletion of γ1-COP or γ2-COP affects Golgi morphology. (a) Electron micrographs of thin sections obtained from P19 WT, *Copg1*^-/-^ and *Copg2*^-/-^ cells as indicated. Identified Golgi stacks are indicated by a yellow G. Quantification of (b) average number of Golgi stacks per cell, (c) average area per Golgi stack and (d) percentage of Golgi stacks with more than 50% of vesicles per stack (p-value: * <0.05, ** < 0.01, ns: non-significant, n=30).

### Depletion of γ1-COP or γ2-COP does not lead to elevated ER stress

ER stress is known to induce Golgi fragmentation^22^ and mutations within COP subunits may lead to ER stress^12, 23^. As ER stress was reported to affect neuronal differentiation of P19 cells^24^, we investigated if depletion of γ1-COP or γ2-COP induces ER stress by assessing expression levels of the ER chaperone GRP-78 (alternatively known as Bip), a commonly used ER stress marker^25, 26^. After western blot analysis, no obvious difference in GRP-78 expression was observed between WT, *Copg1^-/-^* and *Copg2^-/-^* cells (Fig. 4a). This is in contrast to WT cells in which ER stress was induced by tunicamycin, a glycosylation inhibitor that induces the unfolded protein response^27^. Altogether our data suggest that depletion of γ1-COP or γ2-COP does not lead to elevated ER stress.

**Figure 4:**
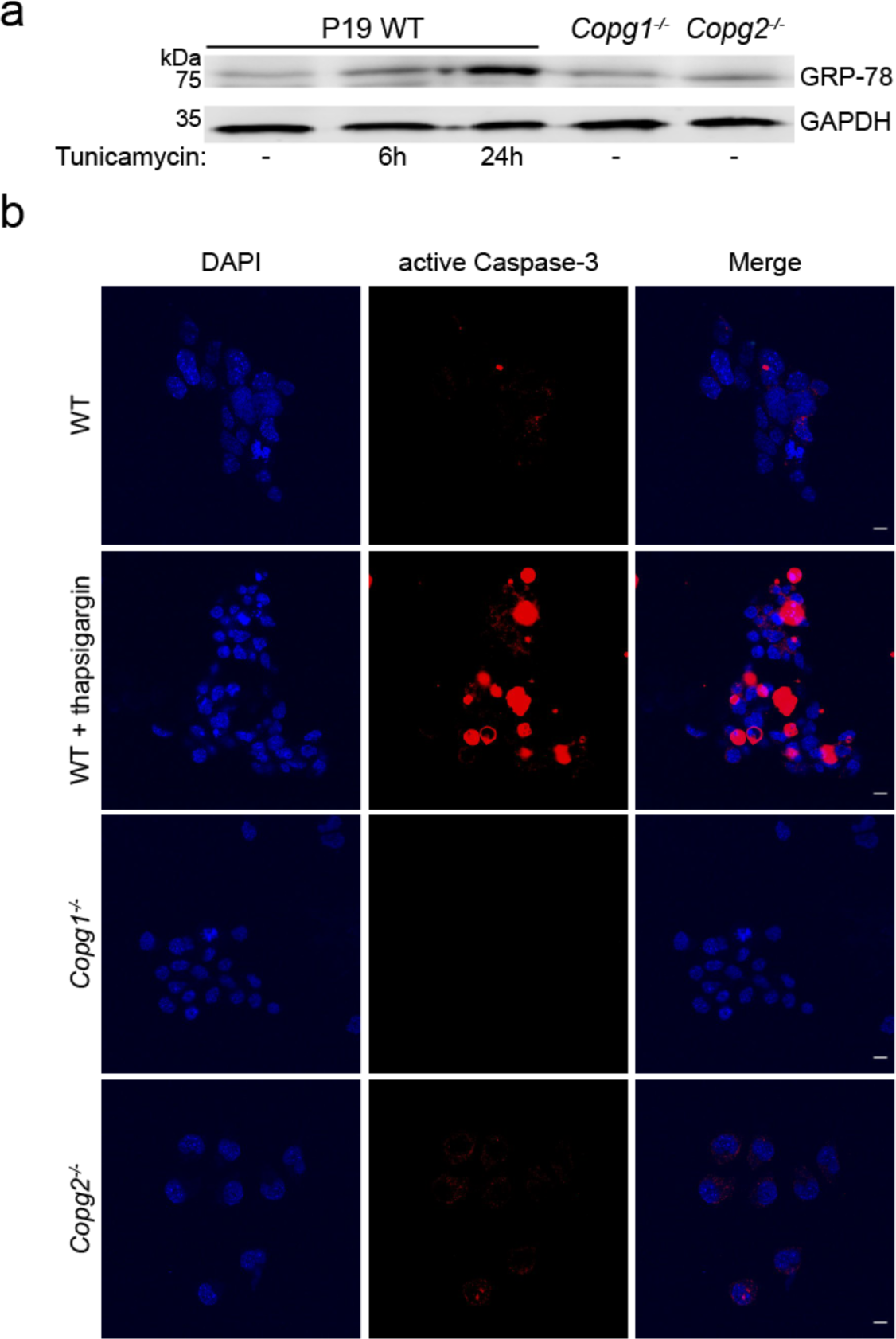
Outcome of γ1-COP or γ2-COP depletion on ER stress and apoptosis. (a) Western blot analysis of the expression level of the ER stress marker GRP-78 and the loading control GAPDH in P19 WT, *Copg1*^-/-^ and *Copg2*^-/-^ cells. ER stress was induced in WT cells with tunicamycin as indicated. (b) Immunostaining of the apoptosis marker cleaved Caspase-3 in P19 WT, *Copg1*^-/-^ and *Copg2*^-/-^ cells. Apoptosis was induced in WT cells with thapsigardin as indicated. Scale bar: 10 μm.

### Depletion of γ2-COP leads to detectable increase of apoptosis

Golgi fragmentation can also be a consequence of apoptosis^28^. Since depletion of γ1-COP or γ2-COP led to reduced proliferation rates (Fig. 2d), we asked if both proteins are needed to maintain P19 cells survival. To do so, cells were visualized by indirect immuno-fluorescence microscopy with an antibody against the active form of Caspase-3, a marker for apoptosis. Many apoptotic cells were observed when P19 WT cells were treated with thapsigargin, an ER stress-inducing drug that leads to apoptosis^29^. Only occasional active Caspase-3 staining was observed in non-treated P19 WT and *Copg1^-/-^* cells (Fig. 4b). By contrast, slightly more *Copg2^-/-^* cells showed active Caspase-3 staining, indicating increased apoptosis upon γ2-COP depletion.

### Expression of γ1-COP is necessary for the formation of tight embryonic bodies

We went on to analyze whether γ-COP paralogs have specific functions during neurogenesis. To address this, P19 WT and KO cells were submitted to the two-step differentiation protocol described in Fig. 1a, in which cells are first led to aggregate under non-adherent conditions to form EBs. In initial experiments we performed EB formation in bacterial dishes as described in the original differentiation protocol^30^. Under these conditions we noticed that EBs generated from *Copg1^-/-^* cells seemed smaller and exhibited a less compact morphology than EBs from WT and *Copg2^-/-^* cells. However, as EBs were free floating in the dishes it was difficult to obtain pictures and to perform quantitative analyses. To investigate EB morphology in a more controlled way we turned to a hanging drop protocol in which 200 dissociated cells are seeded in hanging drops of 20 μL RA-containing cell culture medium. The drops were then analyzed by light microscopy at day 2 and day 4 of differentiation. Typically, in the WT situation, virtually all cells had aggregated to form a single, sometimes two, spherical EBs with sharp boundaries per drop (Fig. 5a). At day 4, EBs appeared slightly larger and denser than at day 2 and also sometimes started to disaggregate. The morphology of EBs obtained from *Copg2^-/-^* cells was similar to WT EBs indicating that γ2-COP is not necessary for EB formation (Fig. 5b). In contrast, EBs obtained from *Copg1^-/-^* cells had a strikingly distinct morphology with much less well-defined shapes and boundaries, and a general looser and more scattered appearance. At day 4 of differentiation, the difference was even more striking with hardly recognizable tight EBs and many disaggregated cells in the drops (**Fig. 5c**).

**Figure 5:**
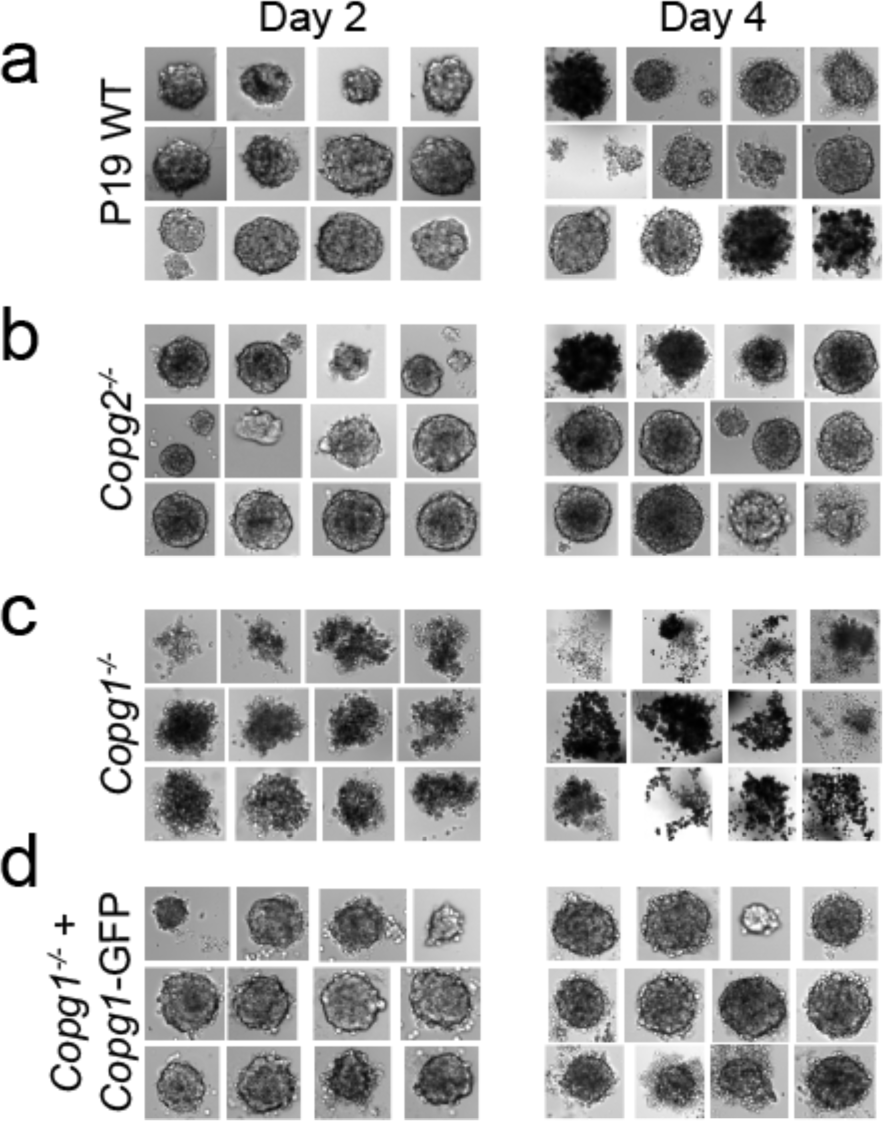
Depletion of γ1-COP affects the formation of embryoid bodies. Photographs of EBs from P19 WT (a), *Copg2*^-/-^ (b), *Copg1*^-/-^ (c), and *Copg1*^-/-^ + *Copg1*-GFP (d) cells formed in hanging drops over 2 or 4 days of culture as indicated.

These observations suggest that expression of γ1-COP is necessary for EB formation. However, as the cell line used in these experiments was obtained from a clonal selection following a Cas9-induced double-strand break within the *Copg1* gene, we decided to perform additional experiments aimed at addressing potential off-target effects of the sgRNA or clone-specific phenotypes. First, we constructed another *Copg1* KO cell line using a different strategy. Instead of inducing the formation of indels within the *Copg1* gene, the whole *Copg1* locus was removed by two single cuts induced by the Cas12a enzyme guided by two crRNAs, and replaced by the coding sequence of either GFP (P19 GFP KI cell line) or of γ1-COP-GFP (P19 γ1-COP-GFP KI cell line) as a control (Supplementary Fig. 3a, b, and see methods). In these knock-in (KI) cell lines, disruption of the *Copg1* locus (GFP KI) also led to strongly impaired EB formation, whereas replacement of the *Copg1* locus by the coding sequence of γ1-COP-GFP had no effect (Supplementary Fig. 3c). The fact that both the *Copg1^-/-^* and GFP KI cell lines show similar phenotypes but are two different clones derived from two different genome editing strategies strongly suggests that the absence of γ1-COP is the cause of defective EB formation. As a further specificity control, we followed a rescue approach in which γ1-COP was re-introduced into *Copg1^-/-^* cells. We first constructed rescued cell lines through the transfection of *Copg1* and *Copg2* KO cells with a PiggyBac (PB) transposon vector driving the constitutive expression of γ1-COP and γ2-COP, respectively, through a strong CAG promoter (resulting in *Copg1^-/-^*-PB-*Copg1* and *Copg2^-/-^*-PB-*Copg2* cell lines). However, western blot analysis revealed that strong overexpression of one paralog led to a much weaker expression of the other paralog (Supplementary Fig. 4a). In fact, *Copg1^-/-^*-PB-*Copg1* cells appeared like a mimic of *Copg2* KO cells whereas *Copg2^-/-^*-PB-*Copg2* cells looked like a mimic of *Copg1* KO cells (Supplementary Fig. 4a, compare lane 3 to 5 and 6 to 2). Previous studies showed that, except for ζ-COP, individual COP subunits are not detectable as monomers in the cytoplasm and only exist as part of the heptameric coatomer complex^31, 32^. We therefore assume that the overexpressed γ-COP efficiently competes with its endogenous counterpart for assembly in the coatomer complex and that the excessive non-assembled protein gets rapidly degraded. As our goal was to obtain a rescue cell line with both γ-COP paralogs expressed close to their endogenous concentrations, we then turned to another strategy in which we used a bacterial artificial chromosome (BAC) that carries the whole *Copg1* gene locus under the control of its native promoter. Through recombineering the *Copg1* gene was fused at the 3’ end to the localization and affinity purification (LAP) tag that comprises a GFP sequence^33^. This strategy ensures that the re-introduced transgene is expressed at levels similar to the endogenous gene and under the control of its native regulatory elements^34^. Western blot analysis of lysates of the rescue cell line (termed P19 *Copg1^-/-^ + Copg1*-GFP) revealed that γ1-COP-GFP was indeed expressed at similar levels to endogenous γ1-COP in the WT cell line (Supplementary Fig. 4a, lane 4). Moreover, by disrupting *Copg2* in this rescued cell line, we successfully obtained P19 cells that solely express γ1-COP-GFP in a double *Copg1^-/-^, Copg2^-/-^* background (Supplementary Fig. 4b). As the COPI pathway is essential, this demonstrates that GFP-tagged γ1-COP is functional. Accordingly, the P19 *Copg1^-/-^ + Copg1*-GFP rescued cell line showed faster proliferation rates than *Copg1^-/-^* cells (Supplementary Fig. 4c) and formation of tight EBs (Fig. 5d). Hence, altogether, our data show that it is indeed the absence of the γ1-COP protein that prevents proper formation of EBs during neuronal differentiation of *Copg1* KO cells.

### Expression of γ1-COP promotes neurite outgrowth

We then analyzed WT and KO cells during the second step of neuronal differentiation, in which EB cells are dissociated, seeded on poly-lysine-coated plates (day 5 of differentiation) and grown for an additional four days (until day 8 of differentiation). Two days after plating, cytosine arabinoside (ara-C) was added to poison dividing cells and hence enrich for post-mitotic neurons. During the second stage, P19 WT cells efficiently differentiated into neurons with the strong upregulation of βIII-tubulin and the extension of long neurites (Fig. 6a, top row). P19 *Copg2^-/-^* cells showed the same hallmarks of neuronal differentiation as WT cells with increased βIII-tubulin expression and long neurites (Fig. 6a, second row from top). By contrast, *Copg1^-/-^* cells hardly showed any long neurites at day 8 of differentiation even though the cells expressed βIII-tubulin (Fig. 6a, third row from top). The P19 *Copg1^-/-^ + Copg1*-GFP rescued cell line showed similar neurite extensions to WT and *Copg2^-/-^* cells (Fig. 6a, bottom row). Further demonstrating the specificity of the *Copg1^-/-^* cell phenotype, the GFP KI cell line also hardly showed any extended neurites whereas the γ1-COP-GFP KI cell line behaved similar to the WT cells (Supplementary Fig. 5). Semi-supervised automated measurements of neurite lengths from WT, KO and rescued cells were performed using the NeuriteQuant software^35^. The quantification corroborated the qualitative observations with a similar higher average neurite length per cell for WT, *Copg2^-/-^* and rescued *Copg1^-/-^* cells compared to *Copg1^-/-^* cells (Fig. 6b).

**Figure 6:**
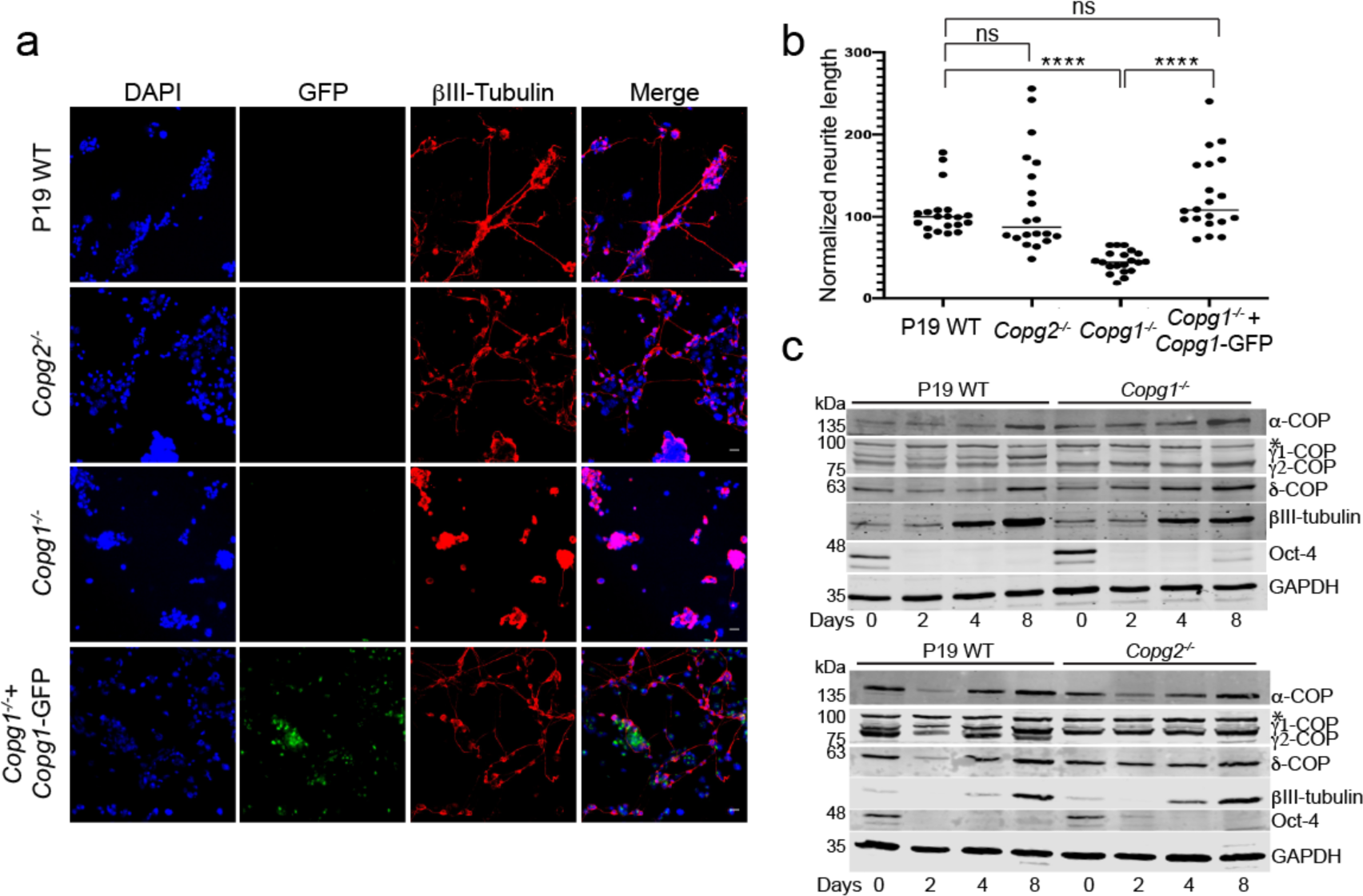
Depletion of γ1-COP leads to impaired neurite outgrowth. (a) Representative fluorescence microscopy images of P19 WT, *Copg2*^-/-^, *Copg1*^-/-^, and *Copg1*^-/-^ + *Copg1*-GFP cells as indicated at day 8 of differentiation to analyze the expression of the neuronal marker βIII-tubulin (indirect immuno-fluorescence) and GFP (direct fluorescence). (b) Quantification of the normalized neurite length (number of pixel/cell). Each dot represents the value obtained for one picture (usually corresponding to ca. 100 cells) as in (a). 20 random images (a total of ca. 2000 cells) were taken for the analysis. The horizontal bars represent the median value (set to 100 for WT) obtained for each cell line. *** indicates p-value < 0.0001, n=20. Scale bar is 30 μm. (c) Western blot analysis of various COP subunits, βIII-tubulin, Oct-4 and GAPDH in P19 WT, *Copg1*^-/-^ and *Copg2*^-/-^ cells at different time point of differentiation as indicated. The asterisk (*) marks a non-specific signal.

When performing these assays, we reproducibly observed fewer remaining cells on the dishes at day 8 of differentiation when using *Copg1^-/-^* or GFP KI cells, suggesting that these cells differentiate less efficiently. Indeed, if the γ1-COP-lacking cells yielded fewer post-mitotic neurons they would be more sensitive to the Ara-C treatment, which is toxic to dividing cells. To explore this possibility, we performed the differentiation protocol in the presence or absence of Ara-C for P19 WT, *Copg1^-/-^* and GFP KI cell lines. For each cell line we then analyzed the percentage of recovered cells after Ara-C treatment (Supplementary Fig. 6a). We found that both γ1-COP-lacking cell lines were more sensitive to Ara-C than WT cells, indirectly suggesting that they differentiate less efficiently into post-mitotic neurons. In support to this hypothesis, immunofluorescence microscopy analysis at day 8 of differentiation showed much less cells positive for the neuronal marker Tuj-1 in the absence of Ara-C treatment for *Copg1^-/-^* and GFP KI cells than in the WT situation (Supplementary Fig. 6d). Importantly, *Copg1^-/-^* and GFP KI cells did not completely fail to differentiate. Indeed, similar to WT and *Copg2^-/-^* cells, *Copg1^-/-^* cells rapidly lost expression of Oct-4 (Fig. 6c) indicating loss of pluripotency. Moreover, the remaining *Copg1^-/-^* and GFP KI cells after Ara-C treatment showed expression of βIII-tubulin (Fig. 6a, c and Supplementary Fig. 5a), implying a correctly implemented neuronal differentiation program.

Altogether the data suggest that expression of γ1-COP regulates the efficiency of neuronal differentiation and promotes neurite outgrowth.

### Generation of P19 cells expressing γ2-COP from the *Copg1* Locus

In non-differentiated *Copg1^-/-^* cells the absence of γ1-COP is at least partly compensated by an increased expression of γ2-COP that seems to maintain global coatomer levels (Fig. 2a-c). However, as we observed an increase of γ1-COP expression at the expense of γ2-COP in neurons derived from P19 WT cells (Fig. 1), it is possible that *Copg1^-/-^* cells do not express enough γ2-COP to compensate at the neuronal stage, thereby restricting the amount of assembled coatomer, and possibly leading to the impaired neurite outgrowth phenotype. To investigate this possibility, we first analyzed the expression levels of α-, γ1-, γ2 and δ-COP in WT, and *Copg1* and *Copg2* KO cells at the non-differentiated, EB and neuron stages (Fig. 6c). In contrast to WT cells, *Copg1^-/-^* cells showed an increased expression of γ2-COP at the neuron stage, while expression levels of the other COP subunits seemed comparable in all three cell lines at all the analyzed differentiation stages. However, as previous estimations found that there is about twice as much γ1-COP than γ2-COP in various cell lines^19, 36^, it is possible that the increased expression of γ2-COP in *Copg1^-/-^* cells is not sufficient to maintain physiological levels of γ-COP in P19 cells throughout differentiation. In that case, we cannot exclude that in *Copg1^-/-^* cells a substantial portion of the cellular coatomer simply misses the γ-COP subunit, as was previously suggested in β-COP-depleted cells^37^.

To characterize the capacity of γ2-COP to replace γ1-COP in a more defined way, we decided to substitute the γ2-COP coding sequence for γ1-COP at the endogenous *Copg1* locus. We thus generated a P19 γ2-COP-GFP KI cell line following the same Cas12a-mediated genome editing strategy that was used to generate the P19 GFP KI and P19 γ1-COP-GFP KI cell lines (Supplementary Fig. 3a, b).

### Expression of γ2-COP from the *Copg1* locus or overexpression of γ2-COP rescues the formation of tight EBs

We next assessed if EB formation specifically requires the γ1-COP protein or just the expression of enough γ-COP irrespective of the paralog identity. To do so P19 WT, GFP KI, γ1-COP-GFP KI and γ2-COP-GFP KI cell lines were used to form EBs in hanging drops. As shown above, whereas WT and γ1-COP-GFP KI formed tight EBs, GFP KI cells showed scattered EBs (Fig. 7a). Strikingly, γ2-COP-GFP KI cells were able to form tight EBs similar to WT cells (Fig. 7a, bottom rows).

**Figure 7:**
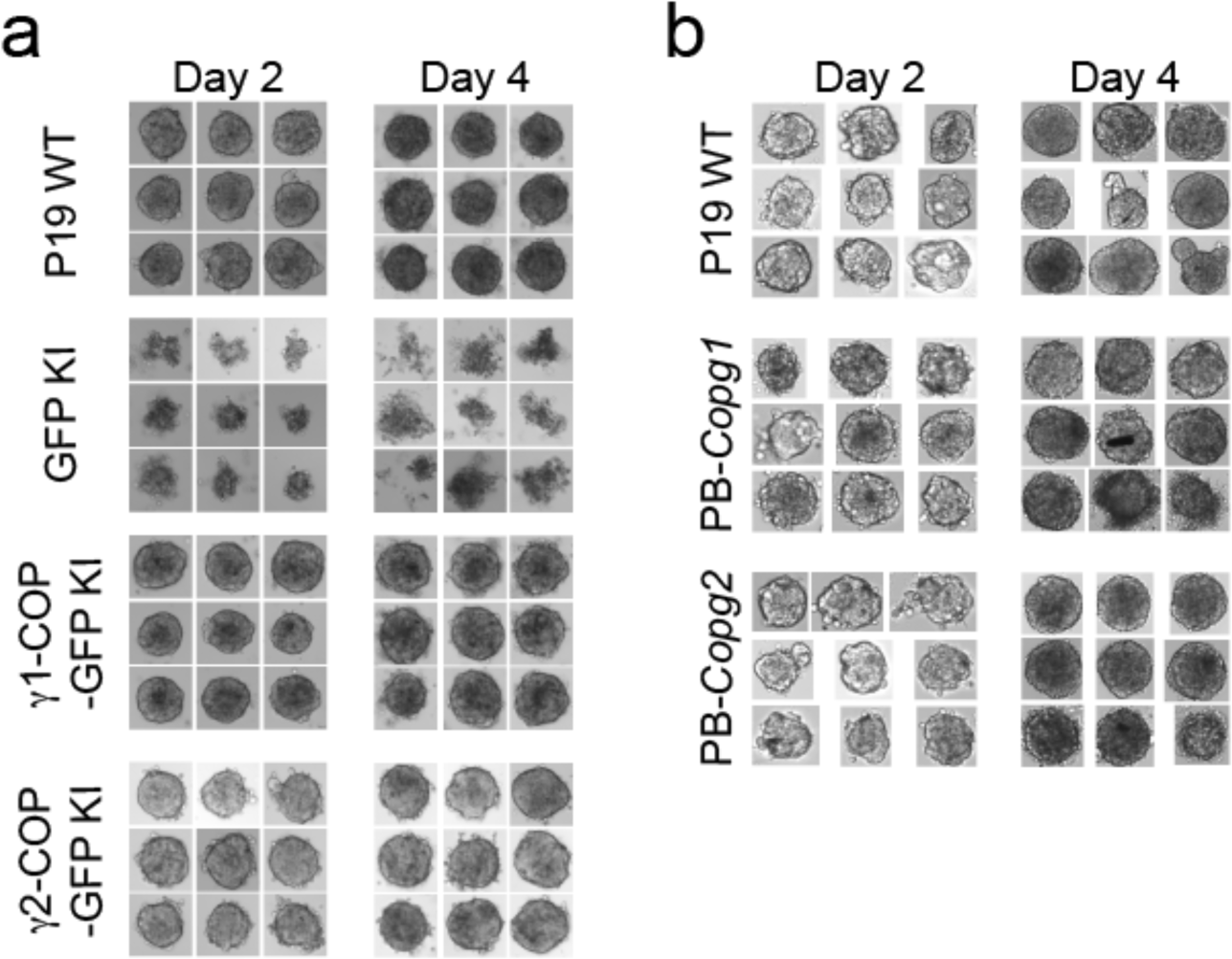
Overexpression of γ2-COP rescues the formation of tight embryoid bodies. (a) Photographs of EBs from P19 WT, GFP-KI, *Copg1*^-/-^ + γ1-COP-GFP KI and *Copg1*^-/-^ + γ2-COP-GFP cells formed in hanging drops over 2 or 4 days of culture as indicated. (b) Same as in (a) with P19 WT, PB-*Copg1* and PB-*Copg2* cells.

Since these cells do not express γ1-COP, this demonstrates that the phenotype of *Copg1^-/-^* and GFP KI cells is not due to a missing specific function mediated by γ1-COP. It rather appears that the absence of γ1-COP is not fully compensated by the upregulation of endogenous γ2-COP in *Copg1^-/-^* cells, but that otherwise both paralogs can support the formation of EBs. To further explore this possibility, we turned to the PiggyBac rescued cell lines we initially generated to rescue *Copg1* and *Copg2* KO cells. Indeed, in these two cell lines, overexpression of one or the other γ-COP paralog leads to essentially one population of coatomer complex that contains the overexpressed protein (Supplementary Fig. 4a). Strikingly both rescue cell lines showed EB formation comparable to WT cells (Fig. 7b). Whereas this was expected for *Copg1^-/-^*-*PB-Copg1* cells as they mimic *Copg2* KO cells, the phenotype of *Copg2^-/-^*-*PB-Copg2* cells suggests that when enough γ2-COP is expressed, EB formation can proceed normally even in the absence of γ1-COP. Thus, our data show that rather than the presence of a specific γ-COP paralog, it is a sufficient expression level of γ-COP that is essential for the formation of EBs in P19 cells.

### Expression of γ2-COP from the *Copg1* locus or overexpression of γ2-COP does not compensate for the absence of γ1-COP during neurite outgrowth

We next analyzed if the absence of γ1-COP can also be compensated by overexpression of γ2-COP during the second step of *in vitro* neuronal differentiation during which neurite outgrowth occurs. As shown above (Supplementary Fig. 5), microscopic analysis revealed that, whereas P19 GFP KI cells fail to extend long neurites, P19 γ1-COP-GFP KI cells show a network of long neurites comparable to WT cells (Fig. 8a). In addition, P19 γ1-COP-GFP KI cells are more resistant to Ara-C treatment than P19 GFP KI cells (Supplementary Fig. 6b). Interestingly, expressing γ2-COP-GFP from the *Copg1* locus also resulted in a resistance to Ara-C comparable to WT cells (Supplementary Fig. 6c), suggesting an increased differentiation efficiency compared to γ1-COP lacking cells. Moreover, P19 γ2-COP-GFP KI cells could extend more and longer neurites than γ1-COP lacking cell lines, however their length and abundance were not comparable to those of WT cells (Fig. 8a). These observations were corroborated by the software-mediated quantification that showed a significantly lower average neurite length per cell for the γ2-COP-GFP KI cells when compared to WT and γ1-COP-GFP KI cells (Fig. 8b).

**Figure 8:**
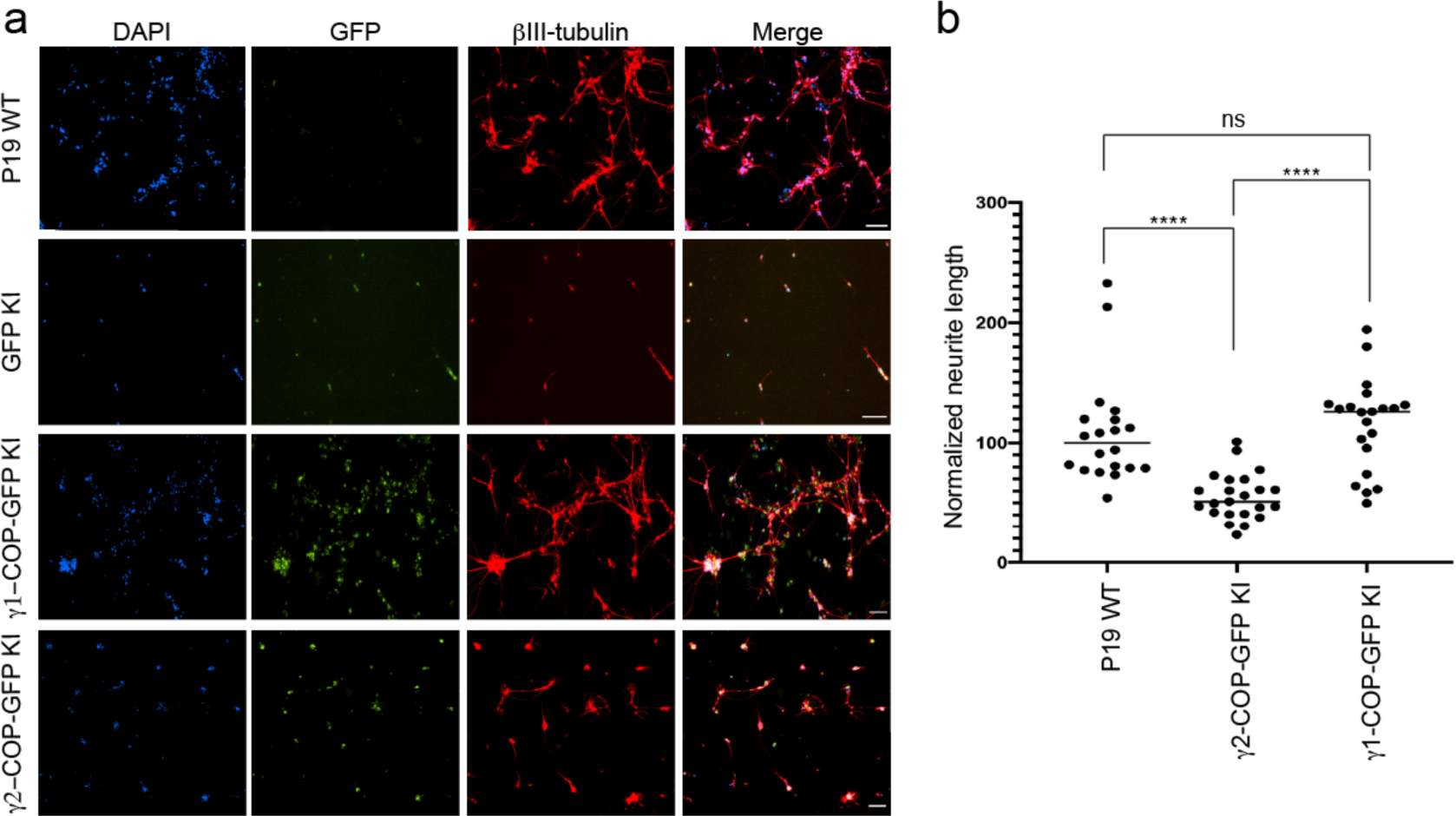
Overexpression of γ2-COP does not compensate the absence of γ1-COP during neurite outgrowth. (a) Representative fluorescence microscopy images of P19 WT, GFP KI, γ1-COP-GFP KI and γ2-COP-GFP KI cells as indicated at day 8 of differentiation to analyze the expression of the neuronal marker βIII-tubulin (indirect immuno-fluorescence) and GFP (direct fluorescence). Scale bar is 100 μm. (b) Quantification of the normalized neurite length (number of pixel/cell). Each dot represents the value obtained for one picture (usually corresponding to ca. 100 cells) as in (a). 20 random images (a total of ca. 2000 cells) were taken for the analysis. The horizontal bars represent the median value (set to 100 for WT) obtained for each cell line. **** indicates p-value < 0.0001, ns: non-significant, n=20.

Hence, in contrast to the EB formation step, it appears that γ1-COP has a specific role during the formation of extended neurites. This was corroborated by the PiggyBac rescued cell lines. Indeed, overexpression of γ1-COP in *Copg1^-/-^* cells (PB-*Copg1* cells) resulted in the formation of neurons with long neurites comparable to the WT situation. In contrast, overexpression of γ2-COP in *Copg2^-/-^* cells (PB-*Copg2* cells) led to the formation of neurons, but with much shorter neurites (Supplementary Fig. 7). Hence, replacement of γ1-COP by an excess of γ2-COP induces a phenotype comparable to the disruption of the *Copg1* gene, suggesting that γ1-COP has a unique role during neuronal differentiation that cannot be taken over by γ2-COP. Of note, as *Copg2^-/-^*-PB-*Copg2* cells form normal EBs but fail to extend long neurites (Fig. 7b), the defective neurite outgrowth observed in *Copg1* KO cells is unlikely to be a mere consequence of their incapacity to form proper EBs.

Altogether, our data reveal a paralog-specific function of the COPI pathway during the neuronal differentiation of P19 cells. In this context, γ1-COP promotes efficient neurite outgrowth in differentiated neurons.

## DISCUSSION

The *Copg2* and *Copz2* genes, paralogous to *Copg1* and *Copz1*, were discovered two decades ago^3, 4^. Since then, it has been an outstanding question whether unique functions can be ascribed to COPI paralogous proteins^38–40^. Recombinant coatomer complexes that contain either γ1-COP or γ2-COP are equally efficient in producing COPI vesicles from purified Golgi membranes^41^ or permeabilized cells^5^. Moreover, the proteomes of COPI vesicles prepared with γ1-COP- or γ2-COP-containing coatomer and using permeabilized HeLa cells as donor membranes were virtually indistinguishable^5^. Hence, these previous *in vitro* data suggested that γ1-COP and γ2-COP are functionally redundant. Our present study performed with living cells revisit this conclusion by showing that whereas γ1-COP and γ2-COP are indeed partially functionally redundant, a paralog-specific function of γ1-COP promotes the extension of neurites in pluripotent cells-derived neurons.

The COPI pathway is essential for life and depletion of non-paralogous COP subunits is lethal in mammalian cells^42, 43^. As we obtained cell lines that lack either γ1-COP or γ2-COP, our data demonstrate that both proteins can support the essential functions of the COPI pathway and that none of the two γ-COP paralogs is essential. However, both *Copg1* and *Copg2* KO cell lines had slower proliferation rates than WT cells indicating that γ1-COP and γ2-COP have some non-essential unique functions. Supporting this hypothesis, whereas disruption of *Copg1* or *Copg2* did not lead to a complete loss of the Golgi as reported upon the depletion of the non-paralogous subunits α- or β-COP^44^, depletion of either γ1-COP or γ2-COP led to smaller and more numerous Golgi stacks.

Rapid evolution of dosage sharing is a main driving force to maintain gene duplicates in mammals^45^. Hence a difficulty when assessing individual knock-out of paralogous gene pairs is to disentangle whether the cause of a phenotype is a unique function that can be assigned to one paralog or an aberrant combined expression level of two paralogous genes with essentially overlapping functions. Indeed, whereas our initial data suggested a role of γ1-COP in promoting the formation of tight EBs, knock-in and overexpression experiments clearly showed that γ2-COP can take over γ1-COP’s role when expressed in sufficient quantity. This suggests that in this context expression of γ1-COP and γ2-COP is regulated by dosage balance to achieve a suitable combined expression of γ-COP compatible with a functional COPI pathway. By contrast, the same combination of experiments supports a unique role for γ1-COP that cannot be taken over by γ2-COP in promoting efficient neurite extension at a later stage of P19 cell neuronal differentiation. This finding correlates with the initial observation that in WT cells, expression of γ1-COP is upregulated at the expense of γ2-COP only at later stages of differentiation, when neurite outgrowth occurs.

What could be the molecular mechanism underlying the role of γ1-COP in promoting neurite outgrowth? Previous studies already hinted at an important role of ER/Golgi trafficking during neuronal polarization. For example, overexpression of the GTP-locked Q71L-ARF1 variant or treatment with the coatomer complex inhibitor 1,3-cyclohexanebis(methylamine)^46^ leads to impaired dendritic growth in cultured hippocampal neurons^47^. Moreover, mutations in the COPII components Sec23 and Sar1 also lead to reduced dendritic growth in flies^48^. In these examples, the outcome of the mutations or treatment is a virtually complete block of ER/Golgi trafficking. This is however not expected in the absence of γ1-COP, especially with the concomitant overexpression of γ2-COP, as γ1-COP-depleted cells are viable and γ2-COP-containing coatomer is as efficient in forming COPI vesicles as γ1-COP-containing coatomer *in vitro*^5, 41^. One possibility is that γ1-COP acts as a specific cargo receptor for proteins that are important for neuronal polarization. In the absence of γ1-COP such hypothetical cargo proteins may be mislocalized, leading to the phenotype of *Copg1* KO cells. The proteomes of COPI vesicles generated with γ1-COP- or γ2-COP-containing coatomer were reported as virtually identical^5^. However, these vesicles were generated from HeLa cells and putative γ1-COP-specific cargo proteins might be neuron-specific. The γ-COP paralogs show a differential Golgi localization with γ1-COP being found preferentially at the *cis*-Golgi and γ2-COP more at the *trans*-Golgi^19^. This suggests that γ-COP paralogs help maintaining protein concentration gradients across the Golgi stack, with γ1-COP being more active at the ER/*cis*-Golgi interface. Proteomic analysis of COPI vesicles generated from various cell lines revealed that they mainly contain membrane trafficking regulating proteins, such as SNAREs (SNAP [soluble NSF attachment protein] receptor), and glycosylation enzymes^5^. The absence of one γ-COP paralog may then affect the distribution or trafficking kinetics of such proteins across the Golgi, which might explain the phenotype of *Copg1* KO cells. Indeed, mutations in both SNAREs^49^ and glycosylation enzymes^50^ are linked to defective neurite outgrowth and neurological disorders. Notably, within the SNARE protein Membrin, pathogenic mutations that lead to less effective fusion of ER-derived vesicles with the Golgi, result in strongly impaired dendritic growth but do not significantly affect non-neuronal cells^51^. This phenotype, which is similar to the depletion of γ1-COP, highlights the requirement of an intact early secretory pathway for neurite outgrowth. As we observed that depletion of γ1-COP results in a more drastic Golgi fragmentation phenotype than depletion of γ2-COP, the two γ-COP paralogs might affect the distribution of Golgi proteins differently, which might explain why only γ1-COP is required for neurite outgrowth.

Finding the proteins that are mislocalized upon depletion of γ1-COP will be key to understanding the molecular mechanisms underlying γ1-COP’s function during neuronal polarization. This is at present not possible with a direct proteomic analysis of isolated COPI vesicles from neurons, considering the amount of starting material needed and the limit of detection of the assay^5^. More promising might be more global approaches such as dynamic organellar maps^52^ or Hyperplexed Localisation of Organelle Proteins by Isotope Tagging (hyperLOPIT)^53^ in which changes in protein localization upon γ1-COP depletion may be analyzed on the proteome-level.

Altogether we found that whereas both γ-COP paralogs can fulfill essential coatomer functions, specialized paralog-specific functions exist. We reveal here a role for γ1-COP in promoting neurite outgrowth, and it is likely that in other cellular contexts additional paralog-specific functions of COP subunits await to be found. Of note, many cancer cell lines are sensitive to a down-regulation of ζ1-COP because they barely express its paralog ζ2-COP^43, 54, 55^. Potential drugs that would target ζ1-COP have thus been proposed as attractive therapeutic options with supposedly limited side effects because ζ2-COP is expressed in normal cells^55, 56^. Our study calls for caution when following such approaches as ζ1-COP, like γ1-COP, may have important functions that cannot be taken over by its paralog in specific cellular contexts.

## MATERIAL AND METHODS

### Antibodies and reagents

Reagents, plasmids, antibodies and primers used in this study are listed in supplementary tables 1-5.

### Cell Culture

The P19 cell line was obtained from Sigma-Aldrich (Germany) and kept at 37°C with 5% CO_2_ in a humidified incubator. The growth medium was Minimum Essential Medium Eagle (α-MEM) supplemented with 10% FBS (Biowest, S181B), 2 mM glutamine and a mixture of penicillin/streptomycin. All the cell lines were regularly checked for mycoplasma contamination.

### P19 differentiation

P19 cells were differentiated into neurons using a two-step protocol essentially as previously described^30^. First, 10^6^ low passage P19 cells were used to seed a 10 cm bacterial dish containing 10 mL of P19 growth medium supplemented with 0.1μM RA. These non-adherent growth conditions promote cell aggregation and the formation of EBs. After 48 h of aggregation the medium was replaced with fresh RA-supplemented P19 growth medium. After 4 days of aggregation, EBs were centrifuged at 1000 *g* for 5 min and then washed once with serum-free media. Then EBs were dissociated using 2 mL of a trypsin (0.05%) - EDTA (0.02%) solution (Sigma, catalog number T3924) + 50μg/mL DNAseI and incubated for 10 min in a 37°C incubator. Next, 4 mL of P19 growth medium were added to stop the trypsin activity and the cells are collected at 1000 *g* for 5 min. Cell pellets were then resuspended in 5 mL growth medium and passed through a cell strainer (Falcon, 352235). Dissociated cells were used to seed poly-L/D-lysine-coated 6-well plates (7.5 x 10^5^ cells in 2 mL per well) or, for fluorescence microscopy experiments, to seed poly-D-lysine-coated 8-well chambered coverslip (Ibidi, 80826) with 40000 cells per well in 300 μL P19 growth medium. 48h after plating (at day 6 of differentiation), the medium was exchanged for P19 growth medium supplemented with 10 μM Ara-C to poison dividing cells. A list of chemicals and their suppliers is provided in supplementary table 1.

### Generation of the P19 *Copg1^-/-^* and *Copg2^-/-^* KO cell lines

P19 *Copg1^-/-^* and *Copg2^-/-^* cells were generated by following a CRISPR-Cas9 gene editing strategy. sgRNAs designed to cut in early exons of the *Copg1* and *Copg2* genes (selected with the CRISPR.MIT tool, see supplementary table 2) were cloned into the pSPCas9(BB)-2TA-GFP vector between the BbsI restriction sites ^57^. P19 cells were transfected with these vectors using Lipofectamine 3000 (Thermo) following manufacturer’s instructions. 72 h after transfection, single GFP-positive cells were sorted and transferred into a 96-well plate using a fluorescence-activated cell sorter (FACS) at the CellNetworks/ZMBH-Flow Cytometry & FACS Core Facility (Heidelberg). Clonal cell lines were grown and transferred into larger cell culture dishes and then screened by western blot. Clones that showed absence of γ1-COP or γ2-COP were verified by pCR-Blunt cloning/Sanger sequencing of the target locus.

### Generation of P19 *Copg1^-/-^*-PB-*Copg1* and *Copg2^-/-^*-PB-*Copg2* cell lines

The P19 *Copg1^-/-^*-PB-*Copg1* and *Copg2^-/-^*-PB-*Copg2* cell lines were generated using the PiggyBac transposon system^58^. P19 *Copg1^-/-^* and *Copg2^-/-^* cells seeded in a 6-well plate were transfected with an equal amount (1.5 μg each) of pPbase plasmid (coding for the PiggyBac transposase) and pCyl50-*Copg1* or pCyl50-*Copg2* plasmid (coding for the gene to be inserted and the hygromycin resistance gene). In addition, a third control cell line was generated by co-transfection of pPbase and a pCyl50-GFP plasmid. 24h after transfection, the cells were transferred to a 10 cm plate. After three days of incubation, hygromycin was added to the growth medium (at 150 μg/ml), and the medium was replenished every 2^nd^-3^rd^ day. After 15 days all GFP-transfected control cells were green as judged by fluorescence microscopy. Transfected PB-*Copg1* and PB-*Copg2* cells were kept on selection media for another 5 days and then analyzed by western blot to check expression of γ1-COP or γ2-COP respectively. Supplementary table 3 lists the plasmids used in this study.

### Generation of the P19 *Copg1^-/-^*-*Copg1*-GFP cell line

BACs harboring the murine *Copg1* locus were GFP-tagged by recombineering as described previously^34^. BAC DNA was isolated from *E. coli* DH10 cells using a BAC prep kit (Macherey-Nagel). P19 *Copg1^-/-^* cells were transfected with 1µg BAC DNA using Effectene (QIAGEN), and stable clonal cell lines were obtained after selection with 500 µg/ml geneticin (Gibco) and FACS sorting.

### Generation of P19 knock-in cells

P19 knock-in cells were generated by following a CRISPR-Cas12a/Cpf1 gene editing strategy including crRNAs arrays, a donor template for HDR (homology directed repair) and a recombination reporter plasmid for selection. Two crRNAs (see supplementary table 2) targeting the *Copg1* gene within the 5’UTR and 3’UTR coding sequencing were selected with the CHOPCHOP tool ^59^ and cloned as a dual array between the BsmbI sites of pY109 (lenti-LbCpf1) plasmid^60^ derivative (pY109_2) in which the Puro-selection cassette was removed. The donor template cassettes were PCR products containing homology arms (left 75bp and right 55 bp for homology directed repair) to the target locus flanking the coding sequence for either GFP (GFP KI cells), γ1-COP-GFP (γ1-COP-GFP KI cells) or γ2-COP-GFP (γ2-COP-GFP KI cells). The donor cassettes were assembled using the NEBuilder HiFi DNA Assembly Master Mix (NEB) and were designed to leave the 5’UTR and 3’UTR of the *Copg1* gene intact. The recombination reporter plasmid (pMB1610_pRR-Puro) allows a split puromycin resistance approach for selecting cells in which Cas12a cut and HDR occurred^61^. The target sequence for crRNA2 was cloned between the two puromycin fragments cassettes within pMB1610. When the Cas12a-induced DNA strand break in the plasmid is repaired via HDR this results in a functional puromycin resistance. P19 cells were seeded at 200,000 cells per well of a 6-well plate. On the next day they were transfected with a mixture of 1.5 μg of modified pY109-crRNA array plasmid, 1 μg of PCR donor cassette and 0.5 μg of pMB1610 reporter plasmid using the JetPrime reagent (Polyplus-transfection) following the manufacturer’s instructions. Puromycin was added 24h later to 4 μg.mL^-1^ in the growth medium and the cells incubated for another 48h. Surviving cells were then transferred to a 10 cm dish and allowed to grow to 70-80% confluence. A pool of GFP-positive cells was then selected by FACS and transferred to a 10 cm dish. The cells were allowed to grow to 70-80% confluence and then single GFP-positive cells were selected and transferred to 96-well plates by FACS. Clonal cell lines were grown and transferred to larger cell culture dishes. Cells that showed a Golgi-like GFP signal by fluorescence microscopy were kept for further screening. Suitable clones were selected after western blot analysis of γ1-COP and γ2-COP expression (absence of endogenous γ1-COP and presence of either γ1-COP-GFP or γ2-COP-GFP). The molecular nature of all genome-edited alleles was verified by PCR (outside homology arms to verify a correct junction of the homology arms with the transgenes, and between exon 8 and 9 of *Copg1* to verify the absence of the endogenous *Copg1* locus) and by pCR-blunt/Sanger sequencing of the complete modified target locus. With these controls we made sure that all three knock-in cell lines had a bi-allelic insertion of the transgenes.

### Hanging drop assay

P19 cells were diluted to 10000 cells/mL in RA-containing P19 growth medium. 30 drops of 20 μL volume (containing 200 cells) were then deposited inside the lid of a 10 cm bacterial dish filled with 10 mL PBS to avoid evaporation. On the second and fourth day of EB formation, images of individual drops were taken on a Nikon TS100F microscope fitted with a CMOS camera (Imaging source, DMK 23UX174) using 10x air objective.

### Immunostaining

For immunostaining, cells were washed with preheated (37°C) PBS, then fixed with PBS supplemented with 4% formaldehyde for 20 min at 37°C. Fixed cells were washed twice with PBS, and then permeabilized with PBS + 0.25% Triton X-100 at room temperature (RT) for 10 min. Cells were then washed twice with PBS + 2% BSA and blocked with PBS + 10% BSA for 30 min at RT. Incubation with primary antibodies diluted in PBS + 2% BSA was then performed either for 1h at RT or overnight at 4°C. Next, cells were washed three times with PBS + 2% BSA and incubated with the relevant secondary antibody diluted in PBS + 2% BSA for 30 min at RT in the dark. Cells were then washed twice with PBS + 2% BSA and incubated with DAPI (4′,6-diamidino-2-phenylindole diluted at 0.1 μg/mL in PBS) for 10 min at RT in the dark. After three brief and gentle washes, the first two with PBS +2% BSA, the last one with PBS, cells were mounted with Ibidi mounting medium. Images were acquired with a Nikon ECLIPSE Ti2 microscope equipped with an Andor Clara DR or Nikon DS-Qi2 camera. Neurite length was assessed after βIII-tubulin staining by analysis with the NeuriteQuant software^35^. For each image, total neurite length was normalized to the number of nuclei counted with the Fiji software. A list of antibodies and their working dilutions is given in supplementary table 4.

### Chemical treatment experiments

To induced apoptosis, P19 cells were treated with thapsigargin (1 μM) for 6 h. Thereafter, immunostaining was performed (see above). To induce ER stress, P19 cells were treated with tunicamycin (2.5 μg.mL^-1^) for 6 h or 24 h. Thereafter, western blot analysis was performed.

### Real-time cell proliferation assay

To assess cell proliferation kinetics, 100,000 cells were seeded per well of a 6-well plate and incubated in an IncuCyte Zoom live-cell analysis system (Essen Bioscience) within a 37°C incubator. Images were acquired in every 4 h for a period of 72 h. Data analysis was performed with the IncuCyte ZOOM Software.

### RNA isolation/cDNA preparation /RT-qPCR

RNA isolation, cDNA preparation and RT-qPCR were performed exactly as described in Ref.^62^. Primers used for qPCR are listed in supplementary table 5.

### Cytosol preparation from adherent cells

For cytosol preparation two confluent 15 cm cell culture dishes were used. Cells were washed twice with PBS and collected by scrapping in 0.5 mL of Lysis-buffer (25 mM Tris, pH 7.4 +150 mM NaCl +1 mM EDTA + Protease inhibitor cocktail). Scrapped cells were lysed by passing them 20 times (10 strokes) through a 21-gauge needle and then 20 times through a 27-gauge needle on ice. Cell lysates were first cleared at 4°C, 800 *g*, 5 min and then at 4°C, 100’000 *g,* 1h. Protein concentration was estimated with a Bradford assay. Typically, a yield of ca. 1.5 mg protein per 15 cm plate was obtained.

### Coatomer immuno-precipitation from cytosolic preparations

Per IP, 10 μL Protein G magnetic beads (NEB, S1430S) were used. Beads were washed twice with IP buffer (25 mM Tris, pH 7.4 + 150 mM NaCl + 1 mM EDTA + 0.025% (v/v) tween 20). After the second wash, 100 μL of CM1 antibody supernatant was added per 10 μL Protein G beads, the tube was then filled to ca. 1.4 mL with IP buffer and incubated for 1h at RT on a rotating wheel. Then, the beads were washed once with IP buffer (1mL). Freshly prepared cytosol corresponding to 500 μg protein was then added to the beads, if needed the tubes were filled to ca. 1.4 mL with IP buffer containing protease inhibitors. Cytosol and beads were incubated for 1h at RT on a rotating wheel. The beads were then washed three times with IP buffer to remove unbound proteins. Bound proteins were eluted by incubating the beads with 20 μL 3x SDS loading buffer at 70°C for 10 min. Beads were then separated from the eluted material on a magnetic rack and the eluted material transferred to a fresh tube.

### Electron microscopy

Cells in a 6-well plate were fixed at RT for 30 min by adding fixative solution (2.5% glutaraldehyde and 2% sucrose in 50 mM KCl, 2.6 mM MgCl_2_, 2.6 mM CaCl_2_, 50 mM cacodylate buffer, pH 7.2). The cells were then rinsed five times for 2 min at RT with 0.1M cacodylate buffer, pH 7.2 and incubated at 4°C for 40 min with Contrast I solution (2% OsO_4_ in 50 mM cacodylate buffer). The cells were then rinsed five times for 1 min at 4°C and then twice for 5 min at RT with mQ water. Thereafter, the cells were incubated at RT for 30 min in the dark with Contrast II solution (0.5% uranyl acetate in mQ water). Finally, the cells were dehydrated with increasing amounts of ethanol and embedded in epon epoxy resin (Polysciences). Ultrathin sections of 60 nm were contrasted with uranyl acetate and lead citrate using an AC20 automatic contrasting system (Leica) and examined with a Jeol 1010 electron microscope (Jeol Europe). Images were taken from WT, *Copg1^-/-^* and *Copg2^-/-^* cell lines. Random images corresponding to 30 Golgi regions per sample were taken and analyzed using the Fiji software.

### Statistical analysis

Statistical significances were calculated using a two-tailed unpaired Student’s *t*-test with the Prism 8 software (Graphpad).

## ACKNOWLEDGEMENTS

We thank Felix Wieland (Heidelberg) for the kind gifts of plasmids and antibodies (see sup. Data). We thank Ina Poser (Dresden) for BAC tagging, and Menna Ahmed for her help handling the Incucyte device. We thank the CellNetworks EM, FACS and Nikon imaging center facilities for their expert support. We thank F. Wieland and Mandy Jeske for their critical comments on the manuscript. This work was supported by the excellence initiative of the German federal and state governments (DFG-EXC81).

## AUTHOR CONTRIBUTIONS

M.G. performed all main experiments and analyzed the data shown on Fig. 1-2, 5-7 and Sup. Fig.1-2, 4-5. X.Z. generated and analyzed the knock-in cell lines (Fig. 7-8; Supp. Fig. 3). M.B. performed the stress (Fig. 4) and cell counting experiments (Sup. Fig. 6). K.L.A.L. performed the apoptosis experiments (Fig. 4). C. H. and J.K. prepared and analyzed the EM samples (Fig. 3). S.S.-D. and M. M.M. generated the BAC-rescued cell line. J.B. designed the study, analyzed the data and wrote the paper. All authors edited the paper.

## REFERENCES

1. Béthune J, Wieland FT. Assembly of COPI and COPII Vesicular Coat Proteins on Membranes. Annu Rev Biophys 47, 63–83 (2018).

2. Barlowe C, Helenius A. Cargo Capture and Bulk Flow in the Early Secretory Pathway. Annu Rev Cell Dev Biol 32, 197–222 (2016).

3. Blagitko N, Schulz U, Schinzel AA, Ropers HH, Kalscheuer VM. gamma2-COP, a novel imprinted gene on chromosome 7q32, defines a new imprinting cluster in the human genome. Hum Mol Genet 8, 2387–2396 (1999).

4. Futatsumori M, Kasai K, Takatsu H, Shin HW, Nakayama K. Identification and characterization of novel isoforms of COP I subunits. Journal of biochemistry 128, 793–801 (2000).

5. Adolf F, et al. Proteomic Profiling of Mammalian COPII and COPI Vesicles. Cell reports 26, 250–265 e255 (2019).

6. Adolf F, Rhiel M, Reckmann I, Wieland FT. Sec24C/D-isoform-specific sorting of the preassembled ER-Golgi Q-SNARE complex. Mol Biol Cell 27, 2697–2707 (2016).

7. Bonnon C, Wendeler MW, Paccaud JP, Hauri HP. Selective export of human GPI-anchored proteins from the endoplasmic reticulum. J Cell Sci 123, 1705–1715 (2010).

8. Mancias JD, Goldberg J. The transport signal on Sec22 for packaging into COPII-coated vesicles is a conformational epitope. Mol Cell 26, 403–414 (2007).

9. Mancias JD, Goldberg J. Structural basis of cargo membrane protein discrimination by the human COPII coat machinery. EMBO J 27, 2918–2928 (2008).

10. Bettayeb K, et al. delta-COP modulates Abeta peptide formation via retrograde trafficking of APP. Proc Natl Acad Sci U S A 113, 5412–5417 (2016).

11. Bettayeb K, et al. Relevance of the COPI complex for Alzheimer’s disease progression in vivo. Proc Natl Acad Sci U S A 113, 5418–5423 (2016).

12. Izumi K, et al. ARCN1 Mutations Cause a Recognizable Craniofacial Syndrome Due to COPI-Mediated Transport Defects. Am J Hum Genet 99, 451–459 (2016).

13. Xu X, et al. Mutation in archain 1, a subunit of COPI coatomer complex, causes diluted coat color and Purkinje cell degeneration. PLoS Genet 6, e1000956 (2010).

14. Li H, Custer SK, Gilson T, Hao le T, Beattie CE, Androphy EJ. alpha-COP binding to the survival motor neuron protein SMN is required for neuronal process outgrowth. Hum Mol Genet 24, 7295–7307 (2015).

15. Tippmann SC, et al. Chromatin measurements reveal contributions of synthesis and decay to steady-state mRNA levels. Molecular systems biology 8, 593 (2012).

16. Kelly GM, Gatie MI. Mechanisms Regulating Stemness and Differentiation in Embryonal Carcinoma Cells. Stem Cells Int 2017, 3684178 (2017).

17. Morassutti DJ, Staines WA, Magnuson DS, Marshall KC, McBurney MW. Murine embryonal carcinoma-derived neurons survive and mature following transplantation into adult rat striatum. Neuroscience 58, 753–763 (1994).

18. Staines WA, Morassutti DJ, Reuhl KR, Ally AI, McBurney MW. Neurons derived from P19 embryonal carcinoma cells have varied morphologies and neurotransmitters. Neuroscience 58, 735–751 (1994).

19. Moelleken J, et al. Differential localization of coatomer complex isoforms within the Golgi apparatus. Proc Natl Acad Sci U S A 104, 4425–4430 (2007).

20. Palmer DJ, Helms JB, Beckers CJ, Orci L, Rothman JE. Binding of coatomer to Golgi membranes requires ADP-ribosylation factor. J Biol Chem 268, 12083–12089 (1993).

21. Lowe M, Kreis TE. In vitro assembly and disassembly of coatomer. J Biol Chem 270, 31364–31371 (1995).

22. Nakagomi S, Barsoum MJ, Bossy-Wetzel E, Sutterlin C, Malhotra V, Lipton SA. A Golgi fragmentation pathway in neurodegeneration. Neurobiol Dis 29, 221–231 (2008).

23. Watkin LB, et al. COPA mutations impair ER-Golgi transport and cause hereditary autoimmune-mediated lung disease and arthritis. Nat Genet 47, 654–660 (2015).

24. Kawada K, et al. Aberrant neuronal differentiation and inhibition of dendrite outgrowth resulting from endoplasmic reticulum stress. J Neurosci Res 92, 1122–1133 (2014).

25. Lee AS. The glucose-regulated proteins: stress induction and clinical applications. Trends Biochem Sci 26, 504–510 (2001).

26. Lee AS. The ER chaperone and signaling regulator GRP78/BiP as a monitor of endoplasmic reticulum stress. Methods 35, 373–381 (2005).

27. Oslowski CM, Urano F. Measuring ER stress and the unfolded protein response using mammalian tissue culture system. Methods in enzymology 490, 71–92 (2011).

28. Hicks SW, Machamer CE. Golgi structure in stress sensing and apoptosis. Biochim Biophys Acta 1744, 406–414 (2005).

29. Dahmer MK. Caspases-2, −3, and −7 are involved in thapsigargin-induced apoptosis of SH-SY5Y neuroblastoma cells. J Neurosci Res 80, 576–583 (2005).

30. Lee JA, Xing Y, Nguyen D, Xie J, Lee CJ, Black DL. Depolarization and CaM kinase IV modulate NMDA receptor splicing through two essential RNA elements. PLoS Biol 5, e40 (2007).

31. Kuge O, et al. zeta-COP, a subunit of coatomer, is required for COP-coated vesicle assembly. J Cell Biol 123, 1727–1734 (1993).

32. Lowe M, Kreis TE. In vivo assembly of coatomer, the COP-I coat precursor. J Biol Chem 271, 30725–30730 (1996).

33. Cheeseman IM, Desai A. A combined approach for the localization and tandem affinity purification of protein complexes from metazoans. Sci STKE 2005, pl1 (2005).

34. Poser I, et al. BAC TransgeneOmics: a high-throughput method for exploration of protein function in mammals. Nat Methods 5, 409–415 (2008).

35. Dehmelt L, Poplawski G, Hwang E, Halpain S. NeuriteQuant: an open source toolkit for high content screens of neuronal morphogenesis. BMC neuroscience 12, 100 (2011).

36. Wegmann D, Hess P, Baier C, Wieland FT, Reinhard C. Novel isotypic gamma/zeta subunits reveal three coatomer complexes in mammals. Mol Cell Biol 24, 1070–1080 (2004).

37. Styers ML, O’Connor AK, Grabski R, Cormet-Boyaka E, Sztul E. Depletion of beta-COP reveals a role for COP-I in compartmentalization of secretory compartments and in biosynthetic transport of caveolin-1. Am J Physiol Cell Physiol 294, C1485–1498 (2008).

38. Arakel EC, Schwappach B. Formation of COPI-coated vesicles at a glance. J Cell Sci 131, (2018).

39. Béthune J, Wieland F, Moelleken J. COPI-mediated transport. J Membr Biol 211, 65–79 (2006).

40. Luo PM, Boyce M. Directing Traffic: Regulation of COPI Transport by Post-translational Modifications. Front Cell Dev Biol 7, 190 (2019).

41. Sahlmuller MC, et al. Recombinant heptameric coatomer complexes: novel tools to study isoform-specific functions. Traffic 12, 682–692 (2011).

42. Hobbie L, Fisher AS, Lee S, Flint A, Krieger M. Isolation of three classes of conditional lethal Chinese hamster ovary cell mutants with temperature-dependent defects in low density lipoprotein receptor stability and intracellular membrane transport. J Biol Chem 269, 20958–20970 (1994).

43. Shtutman M, et al. Tumor-specific silencing of COPZ2 gene encoding coatomer protein complex subunit zeta 2 renders tumor cells dependent on its paralogous gene COPZ1. Proc Natl Acad Sci U S A 108, 12449–12454 (2011).

44. Razi M, Chan EY, Tooze SA. Early endosomes and endosomal coatomer are required for autophagy. J Cell Biol 185, 305–321 (2009).

45. Lan X, Pritchard JK. Coregulation of tandem duplicate genes slows evolution of subfunctionalization in mammals. Science 352, 1009–1013 (2016).

46. Hu T, Kao CY, Hudson RT, Chen A, Draper RK. Inhibition of secretion by 1,3-Cyclohexanebis(methylamine), a dibasic compound that interferes with coatomer function. Mol Biol Cell 10, 921–933 (1999).

47. Horton AC, Racz B, Monson EE, Lin AL, Weinberg RJ, Ehlers MD. Polarized secretory trafficking directs cargo for asymmetric dendrite growth and morphogenesis. Neuron 48, 757–771 (2005).

48. Ye B, Zhang Y, Song W, Younger SH, Jan LY, Jan YN. Growing dendrites and axons differ in their reliance on the secretory pathway. Cell 130, 717–729 (2007).

49. Ulloa F, Cotrufo T, Ricolo D, Soriano E, Araujo SJ. SNARE complex in axonal guidance and neuroregeneration. Neural Regen Res 13, 386–392 (2018).

50. Moll T, Shaw PJ, Cooper-Knock J. Disrupted glycosylation of lipids and proteins is a cause of neurodegeneration. Brain, (2019).

51. Praschberger R, et al. Mutations in Membrin/GOSR2 Reveal Stringent Secretory Pathway Demands of Dendritic Growth and Synaptic Integrity. Cell reports 21, 97–109 (2017).

52. Itzhak DN, Tyanova S, Cox J, Borner GH. Global, quantitative and dynamic mapping of protein subcellular localization. eLife 5, (2016).

53. Christoforou A, et al. A draft map of the mouse pluripotent stem cell spatial proteome. Nat Commun 7, 8992 (2016).

54. Anania M, et al. Identification of thyroid tumor cell vulnerabilities through a siRNA-based functional screening. Oncotarget 6, 34629–34648 (2015).

55. Anania MC, et al. Targeting COPZ1 non-oncogene addiction counteracts the viability of thyroid tumor cells. Cancer Lett 410, 201–211 (2017).

56. Shtutman M, Roninson IB. A subunit of coatomer protein complex offers a novel tumor-specific target through a surprising mechanism. Autophagy 7, 1551–1552 (2011).

57. Ran FA, Hsu PD, Wright J, Agarwala V, Scott DA, Zhang F. Genome engineering using the CRISPR-Cas9 system. Nat Protoc 8, 2281–2308 (2013).

58. Wang W, et al. Chromosomal transposition of PiggyBac in mouse embryonic stem cells. Proc Natl Acad Sci U S A 105, 9290–9295 (2008).

59. Labun K, Montague TG, Krause M, Torres Cleuren YN, Tjeldnes H, Valen E. CHOPCHOP v3: expanding the CRISPR web toolbox beyond genome editing. Nucleic Acids Res 47, W171–W174 (2019).

60. Zetsche B, et al. Multiplex gene editing by CRISPR-Cpf1 using a single crRNA array. Nat Biotechnol 35, 31–34 (2017).

61. Flemr M, Buhler M. Single-Step Generation of Conditional Knockout Mouse Embryonic Stem Cells. Cell reports 12, 709–716 (2015).

62. Amaya Ramirez CC, Hubbe P, Mandel N, Bethune J. 4EHP-independent repression of endogenous mRNAs by the RNA-binding protein GIGYF2. Nucleic Acids Res 46, 5792–5808 (2018).

