## Supplementary figures and tables for "A paralog-specific role of the COPI pathway in the neuronal differentiation of murine pluripotent cells"

7 Supplementary figures

5 Supplementary tables

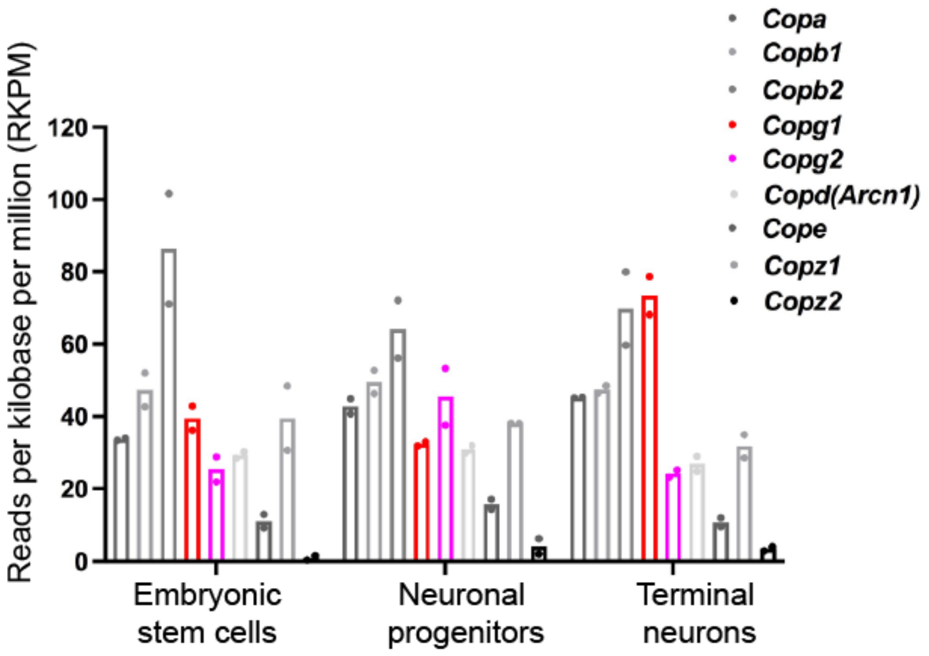


**Supplementary Figure 1: Differential expression of *Copg1* and *Copg2* mRNAs during neuronal differentiation of mouse embryonic stem cells.**
mRNA-seq analysis of cDNAs derived from mouse embryonic stem cells, neuronal progenitors and terminal neurons from two independent differentiation experiments (Tippmann, Ivanek et al. 2012). Normalized RKPM obtained for the COP subunits are indicated. The dataset used to generate this figure is publicly available on the GEO database (accession number GSE34473).

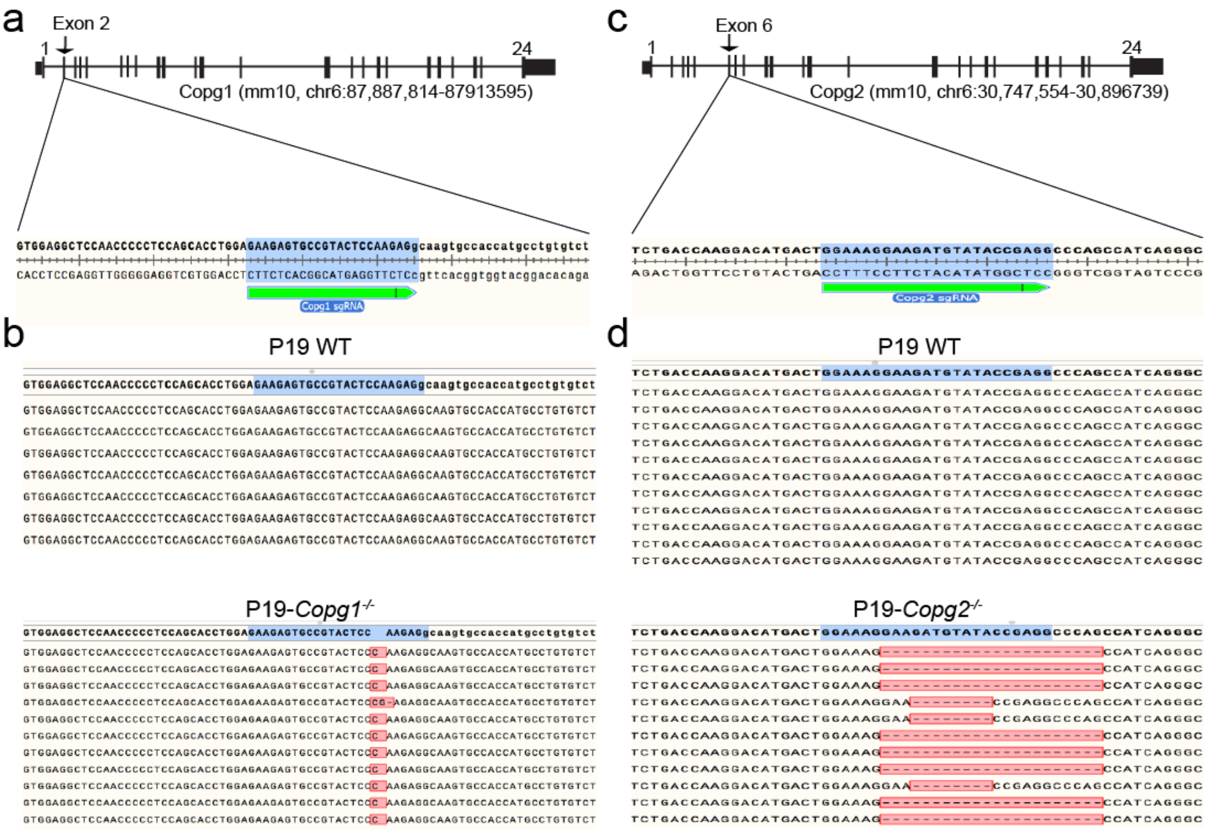


**Supplementary Figure 2: Characterization of P19 *Copg1*$^{-/-}$ and *Copg2*$^{-/-}$ cell lines.**

(a) Schematic of the *Copg1* genomic locus, the sequence targeted by the Copg1 sgRNA is indicated. (b) Sanger sequencing data after PCR amplification of the targeted genomic region, subcloning into a plasmid and screening for individual clones. In bold is the reference sequence. Sequencing from WT cells (top) indicate on intact sequence, sequencing from *Copg1*$^{-/-}$ cells (bottom) indicates two mutated alleles (insertion of one or two nt). (c) same as in (a) with the *Copg2* genomic locus. (d) same as in (c) for the *Copg2* targeted locus in *Copg2*$^{-/-}$ cells. Sequencing indicates two mutated alleles (deletion of 8 or 22 nt).

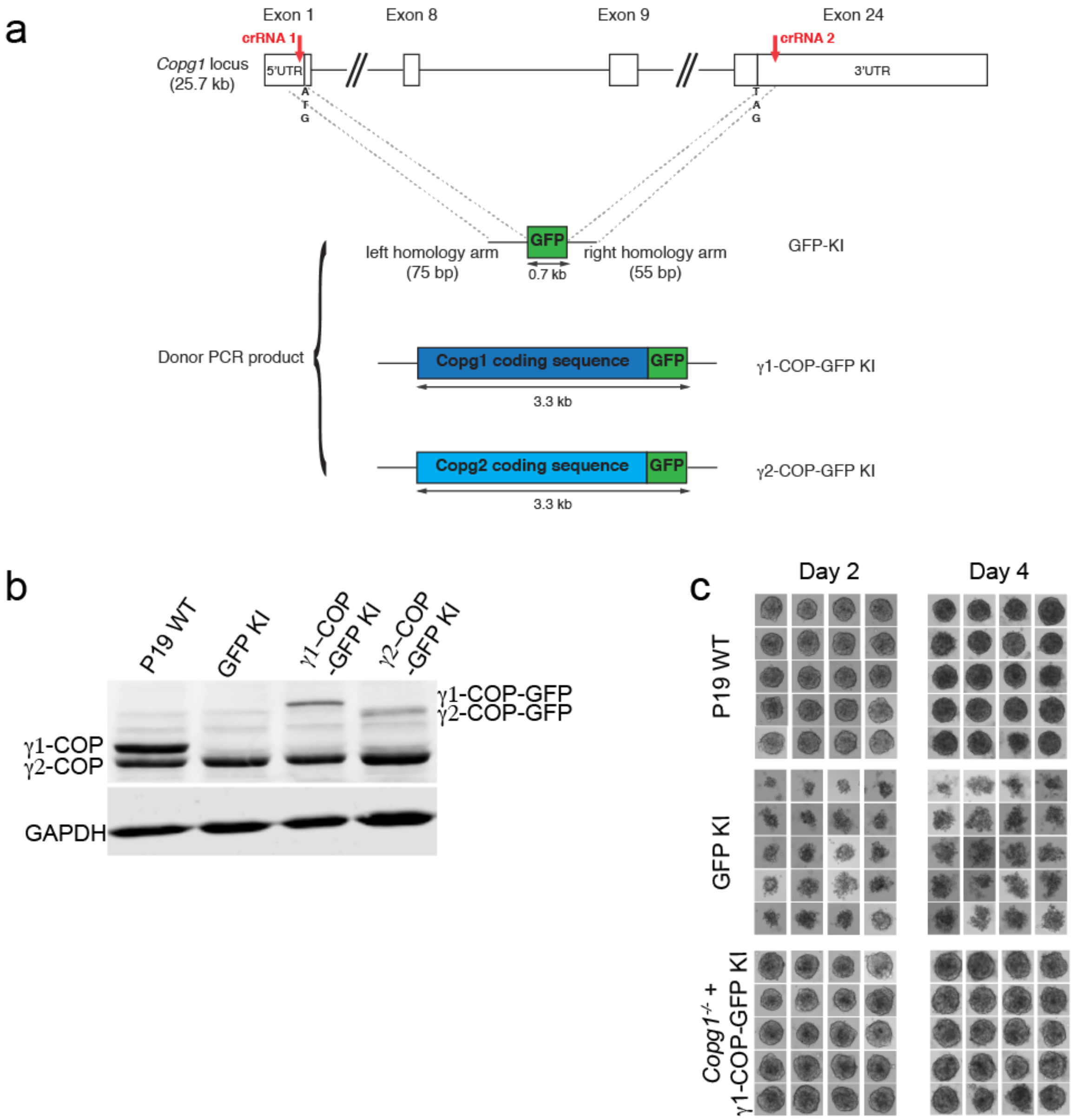


**Supplementary Figure 3: Generation and characterization of P19 knock-in cell lines.**
(a) General strategy to obtain knock-in cell lines. Schematic of the *Copg1* genomic locus, the sites targeted by the crRNAs to guide two Cas12a-induced cleavages are indicated. The three donor PCR products with short homology arms that direct homology directed repair are also represented. (b) Western blot analysis of the expression levels of γ1-COP and γ2-COP, and GAPDH as a loading control in pluripotent P19 WT, GFP KI, γ1-COP-GFP KI and γ2-COP-GFP KI cell lysates. (c) Photographs of EBs from P19 WT, GFP KI and γ1-COP-GFP KI cells formed in hanging drops over 2 or 4 days of culture as indicated.

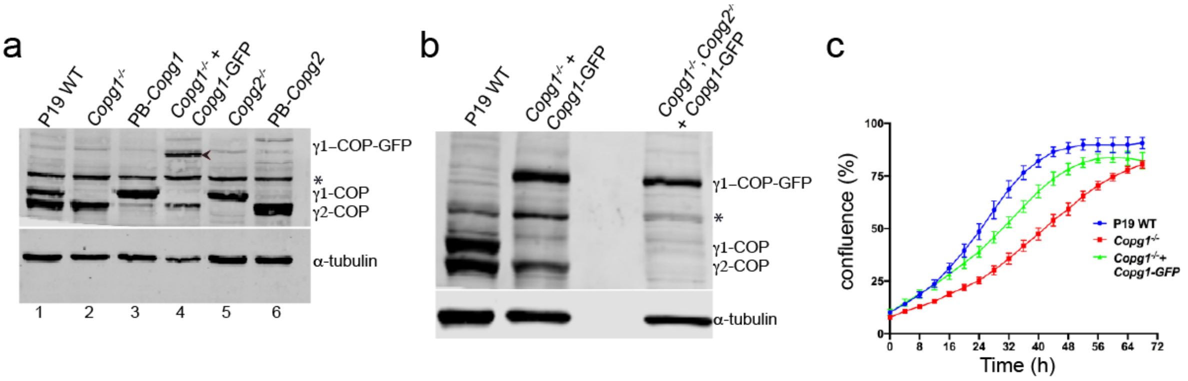


**Supplementary Figure 4: Characterization of rescued P19 cell lines.**
(a) Western blot analysis of the expression levels of γ1-COP and γ2-COP, and α-tubulin as a loading control in pluripotent P19 WT, *Copg1*$^{-/-}$, PB-*Copg1* (*Copg1*$^{-/-}$ cells rescued with a PiggyBac transposon vector), *Copg1*$^{-/-}$ + *Copg1*-GFP (*Copg1*$^{-/-}$ cells rescued with a BAC), *Copg2*$^{-/-}$, PB-*Copg2* cell lysates. The asterisk (*) marks a non-specific signal. The arrow head indicates γ1-COP-GFP in lane 4. (b) Same as in (a) for P19 WT, *Copg1*$^{-/-}$ + *Copg1*-GFP and *Copg1*$^{-/-}$, *Copg2*$^{-/-}$ + *Copg1*-GFP cells. (c) Growth curves obtained from a real-time proliferation assay in which the occupied area (% confluence) by P19 WT, *Copg1*$^{-/-}$ and *Copg1*$^{-/-}$ + *Copg1*-GFP cells was monitored over 72h. Curves were generated with the IncuCyte software (n=5, error bars are s.e.m.).

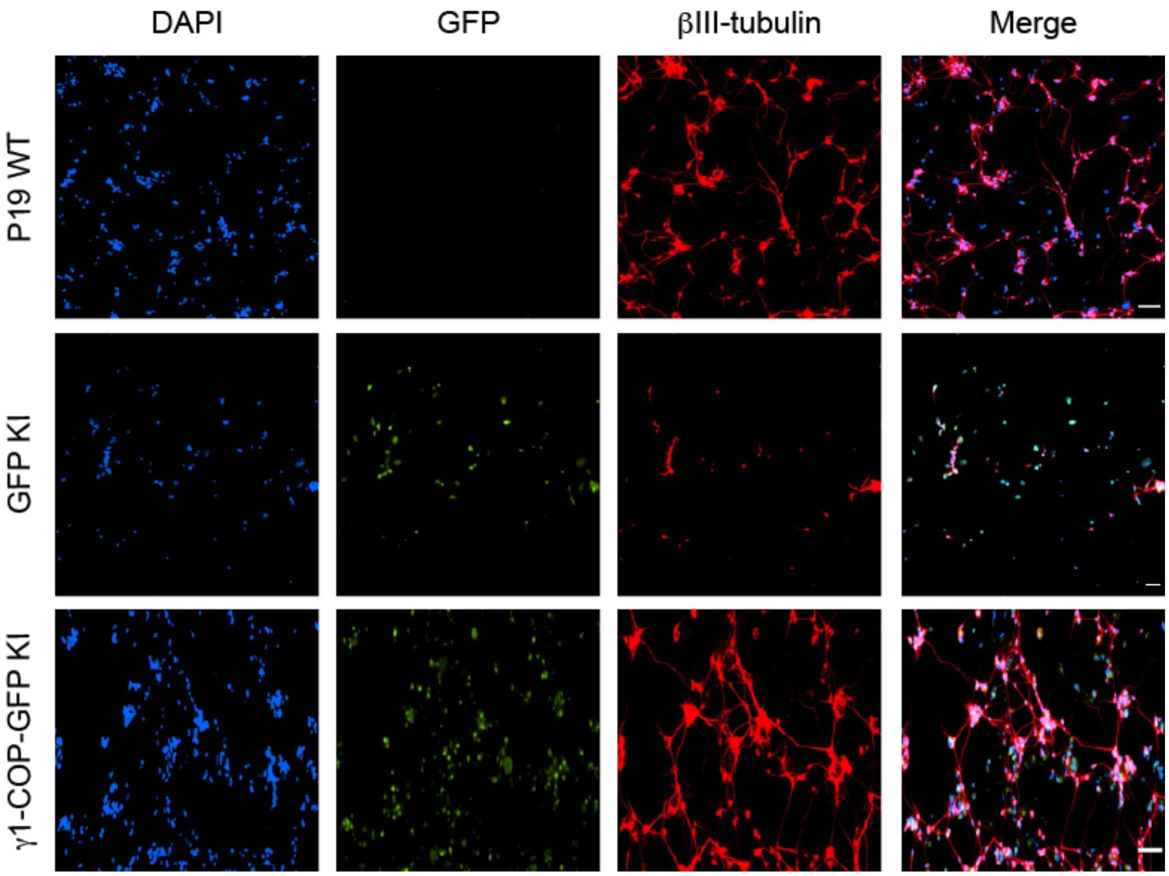


**Supplementary Figure 5: Inhibition of neurite outgrowth in P19 GFP knock-in cells.**

Representative fluorescence microscopy images of P19 WT, GFP KI and γ1-COP-GFP KI as indicated at day 8 of differentiation to analyze the expression of the neuronal marker βIII-tubulin (indirect immuno-fluorescence) and GFP (direct fluorescence). Scale bar is 100 μm.

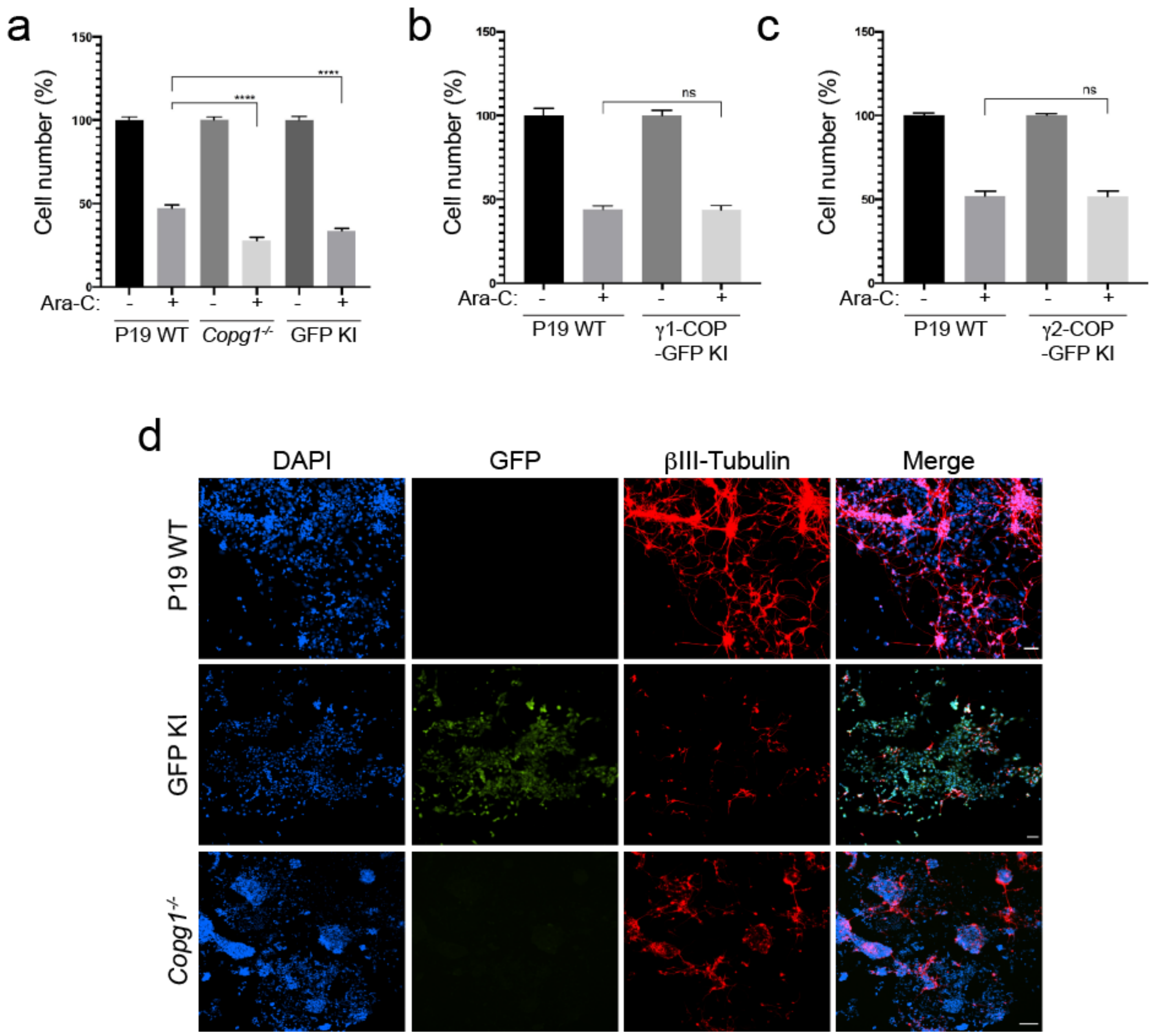


**Supplementary Figure 6: Reduced survival after Ara-C treatment and differentiation efficiency in *Copg1* KO cells.**

(a-c) Quantification of cell survival upon Ara-C treatment. Cells from dissociated EBs were seeded at the same density in a cell culture multi-well plate. Recovered cell numbers after two days of culture with or without Ara-C were calculated. The numbers obtained in the non-treated samples were set to 100%. (a) Comparison between P19 WT, *Copg1*$^{-/-}$ and GFP KI cells. (b) Comparison between P19 WT and γ1-COP-GFP KI cells. (c) Comparison between P19 WT and γ2-COP-GFP KI cells. A two-tailed unpaired t-test was performed for the statistical significance analysis (n=3, **** indicates p-value < 0.0001, ns: non-significant, error bars are s.e.m.). (d) Representative fluorescence microscopy images of P19 WT, GFP KI and *Copg1*$^{-/-}$cells as indicated at day 8 of differentiation without Ara-C treatment to analyze the expression of the neuronal marker βIII-tubulin (indirect immuno-fluorescence) and GFP (direct fluorescence). Scale bar is 100 μm.

a

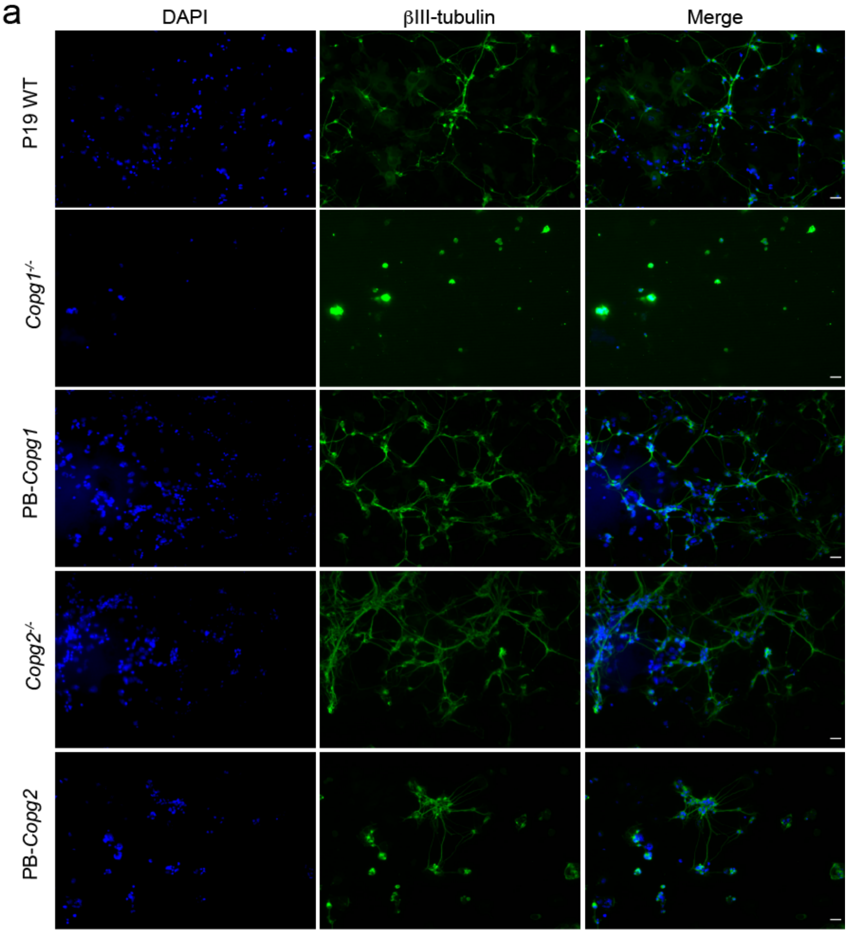


b

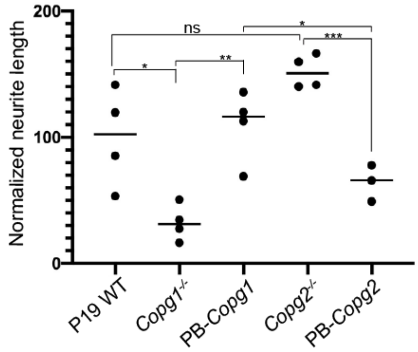


**Supplementary Figure 7: Overexpression of γ2-COP does not compensate the absence of γ1-COP during neurite outgrowth in PiggyBac rescued cells.**
(a) Representative fluorescence microscopy images of P19 WT, *Copg1*$^{-/-}$, PB-*Copg1*, *Copg2*$^{-/-}$, and PB-*Copg2* cells as indicated at day 8 of differentiation to analyze the expression of the neuronal marker βIII-tubulin (indirect immuno-fluorescence). (b) Quantification of the normalized neurite length (number of pixel/cell). Each dot represents the value obtained for one picture (usually corresponding to ca. 100 cells) as in (a). Three to four random images (a total of ca. 400 cells) were taken for the analysis. The horizontal bars represent the median value (set to 100 for WT) obtained for each cell line. A two-tailed unpaired t-test was performed for the statistical significance analysis (n=3 or 4, p-value: * <0.05, ** < 0.01, ***<0.001, n.s: non-significant). Scale bar is 30 µm.

### Supplementary table 1: Materials used in this study

| Chemical | Supplier | Catalog number |
| --- | --- | --- |
| Hygromycin B | Sigma | H0654 |
| Retinoic Acid | Sigma | R2625 |
| poly-D-lysine | Sigma | P0899 |
| Ara-C | Sigma | C1768 |
| Thapsigargin | Caymen Chemicals | Cay10522-1 |
| Tunicamycin | Biomol | Cay11445-10 |
| DNaseI | Applichem | A3778 |
| $\alpha$-MEM | Sigma | M4526 |

### Supplementary table 2: List of sgRNAs/crRNAs

| sgRNA/crRNA | sgRNA sequence | Targeted locus (genomic location, GRCm38) |
| --- | --- | --- |
| *Copg1* sgRNA (cas9) | GAAGAGTGCCGTACTCCAAG | Exon 2 (Ch 6: 87’889’102-121) |
| *Copg2* sgRNA (Cas9) | CGGTATACATCTTCCTTTCC | Exon 6 (Ch 6: 30’863’572-591) |
| *Copg1* crRNA1 (Cas12a) | TTCAACATGGTGTCAGCGGTAGCT | Exon 1 (Ch 6: 87’888’020-043) |
| *Copg1* crRNA2 (Cas12a) | TTAACACAGTGGTCTGGGGAAGGG | Exon 24 (Ch 6: 87’912’378-401) |

### Supplementary table 3: Plasmids used in this study

| Plasmid | Insert | Source |
|---|---|---|
| pSPCas9(BB)-T2A-GFP | Cas9-T2A-GFP | Addgene (48138) |
| pSPCas9(BB)-T2A-GFP *Copg1* sgRNA | Cas9-T2A-GFP + *Copg1* sgRNA | This study |
| pSPCas9(BB)-T2A-GFP *Copg2* sgRNA | Cas9-T2A-GFP + *Copg2* sgRNA | This study |
| pPbase (PL623) | PiggyBac transposase | The Wellcome Trust Sanger Institute |
| pCyl50 | none | The Wellcome Trust Sanger Institute |
| pCyL50-m*Copg1*-Hyg | Mouse $\gamma$1-COP coding sequence | This study |
| pCyL50-m*Copg2*-Hyg | Mouse $\gamma$2-COP coding sequence | This study |
| pY109 (lenti-LbCpf1) | LbCas12a/Cpf1-P2A-puro resistance | Addgene (84747) |
| pY109_2 | LbCas12a/Cpf1 | This study |
| pY109_2-crRNA1/crRNA2 | LbCas12a/Cpf1 + crRNA1/crRNA2 array | This study |
| pMB1610_pRR-Puro | split puromycin N-acetyltransferase coding sequence | Addgene (65853) |

### Supplementary table 4: Antibodies used in this study

| Antibody | Supplier | Application and dilution |
|---|---|---|
| anti-Nanog | Biomol (A300397AT) | WB, 1:1000 |
| anti-Oct-4 | Abcam (ab18976) | WB, 1:500 |
| anti-βIII-Tubulin (TujI/TUBB3) | Biolegend (MMS435P25) | IF, WB, 1:1000 |
| anti-$\alpha$-COP (1409B) | Wieland Lab (Heidelberg) | WB, 1:5000 |
| anti-$\delta$-COP (877) | Wieland Lab (Heidelberg) | WB, 1:1000 |
| anti-$\varepsilon$-COP | Wieland Lab (Heidelberg) | WB, 1:2000 |
| anti-$\gamma$1-COP (anti-$\gamma$1-app) | Wieland Lab (Heidelberg) | WB, 1:500 |
| anti-$\gamma$2-COP (anti-$\gamma$2-app) | Wieland Lab (Heidelberg) | WB, 1:500 |

| | | |
|---|---|---|
| anti-native coatomer (CM1A10/CM1) | Wieland Lab (Heidelberg) | IP: 100 μL of hybridoma supernatant |
| Anti-cleaved Caspase-3 | Cell Signaling (9661T) | IF, 1:100 |
| anti-GAPDH | Proteintech (60004-1-Ig) | WB, 1:20000 |
| anti-α-tubulin | Sigma (T5168) | WB, 1:10000 |
| anti-α-tubulin | Abcam (Ab18251) | WB, 1:10000 |
| Anti-GRP78 | Santa cruz (sc-13539) | WB, 1:200 |

### Supplementary table 5: List of primers used in qPCR experiments

| Gene | IDT prime time Assay Name | Forward primer | Reverse primer |
|---|---|---|---|
| Copa | Mm.PT.58.313 89808 | CAAACCGATTCCGAGCAAC | ACCTACGACCTATACACC ATCC |
| Copb1 | Mm.PT.58.734 1394 | ATAAGCAACATAGCCTCAGCA | CTCGCCACAACTCTAACCAA |
| Copb2 | Mm.PT.58.324 91925 | CCGAAGCTCTTGTTCCTCAA | CCACAGACCATTCAGCACA |
| Copg1 | Mm.PT.58.533 7327 | CTGATGATGCAGTCCACAATG | GTGCCAGAAGTATCCTCGAAAG |
| Copg2 | qMmCID0005 246 (Bio-Rad) | ATCCTACCTCGTTAGCCTGTA | AAGAAGAATGTAAAAGGTGGTGTG |
| Arcn1 | Mm.PT.58.785 9979 | CTCCAAGTTTCAAAGCCTTGC | CAGCCATGATCACAGAGACTATC |
| Copz1 | Mm.PT.58.968 6781 | CCCTCCATCTACAATTTCATCCA | TCTGAACTGCCTCTTCGATTC |
| Copz2 | Mm.PT.58.139 51793 | AAACCATCTGCTCCTTCACG | GAACCTTCTCTCTACACCATCAAG |
| Cope | Mm.PT.58.841 4483 | AGGATCTGAATCGTCATGGC | GACCAATACCACTTTCCTGCT |
